# Unveiling the G4-PAMAM capacity to bind and protect Ang-(1-7) bioactive peptide

**DOI:** 10.1101/2022.05.23.493150

**Authors:** L. América Chi, Somayeh Asgharpour, José Correa-Basurto, Cindy Rodríguez Bandala, Marlet Martínez-Archundia

## Abstract

New therapies that allow natural healing processes are required. Such as the endogenous peptide called Angiotensin-(1-7), a safe and eff e drug, which is able to re-balance the Renin-Angiotensin system affected during several pathologies, including the new COVID-19; cardiovascular, renal, and pulmonary disease; diabetes; neuropathic pain; Alzheimer and cancer. However, one of the limiting factors for its application is its unfavorable pharmacokinetic profile. In this work, we propose the coupling of Angiotensin-(1-7) to PAMAM dendrimers in order to evaluate the capacity of the nanocarrier to improve isolated peptide features and to gain insight into the structural as well as the energetic basis of its molecular binding. The *In Silico* tests were performed in acidic and neutral pH conditions as well as amino-terminated and hydroxyl-terminated PAMAM dendrimers. High-rigor computational approaches, such as molecular dynamics and metadynamics simulations were used. We found that, at neutral pH, PAMAM dendrimers with both terminal types are able to interact stably with 3 Angioteinsin-(1-7) peptides through ASP1, TYR4 and PRO7 key aminoacids, however, there are some differences in the binding sites of the peptides. In general, they bind on the surface in the case of the hydroxyl-terminated compact dendrimer and in the internal zone in the case of the amino-terminated open dendrimer. At acidic pH, PAMAM dendrimers with both terminal groups are still able to interact with peptides either internalized or in its periphery, however, the number of contacts, the percentage of coverage and the number of HBs are lesser than at neutral pH, suggesting a state for peptide release. In summary, amino-terminated PAMAM dendrimer showed slightly better features to bind, load and protect Angiotensin-(1-7) peptides.

## 1 Introduction

The Renin-Angiotensin system (RAS) regulates the blood pressure and also balances ion and other fluids in the human body. For it to function properly, the RAS rests on two opposite axis (Figure 1): 1) in the classical arm, inflammatory, profibrotic, vasoconstrictor and oxidative effects are triggered by the binding of bioactive octapeptide Angiotensin II (Ang-II) to Angiotensin Type 1 (AT1) cell membrane receptor; meanwhile, 2) in the protective arm, opposite anti-inflammatory, anti-fibrotic, vasodilator and anti-oxidant effects are triggered by the binding of the bioactive heptapeptide Angiotensin 1-7 (Ang-(1-7)) to Mitochondrial Assembly (MAS) proto-oncogene which is a G-protein-coupled receptor (GPCRs).^1, 2^ There must be a balance between the two main axes, however, under several pathological conditions this balance is shifted towards the classical arm leading to catastrophic effects, as in the case of the worsening of COVID-19 disease.^3–7^

**Figure 1:**
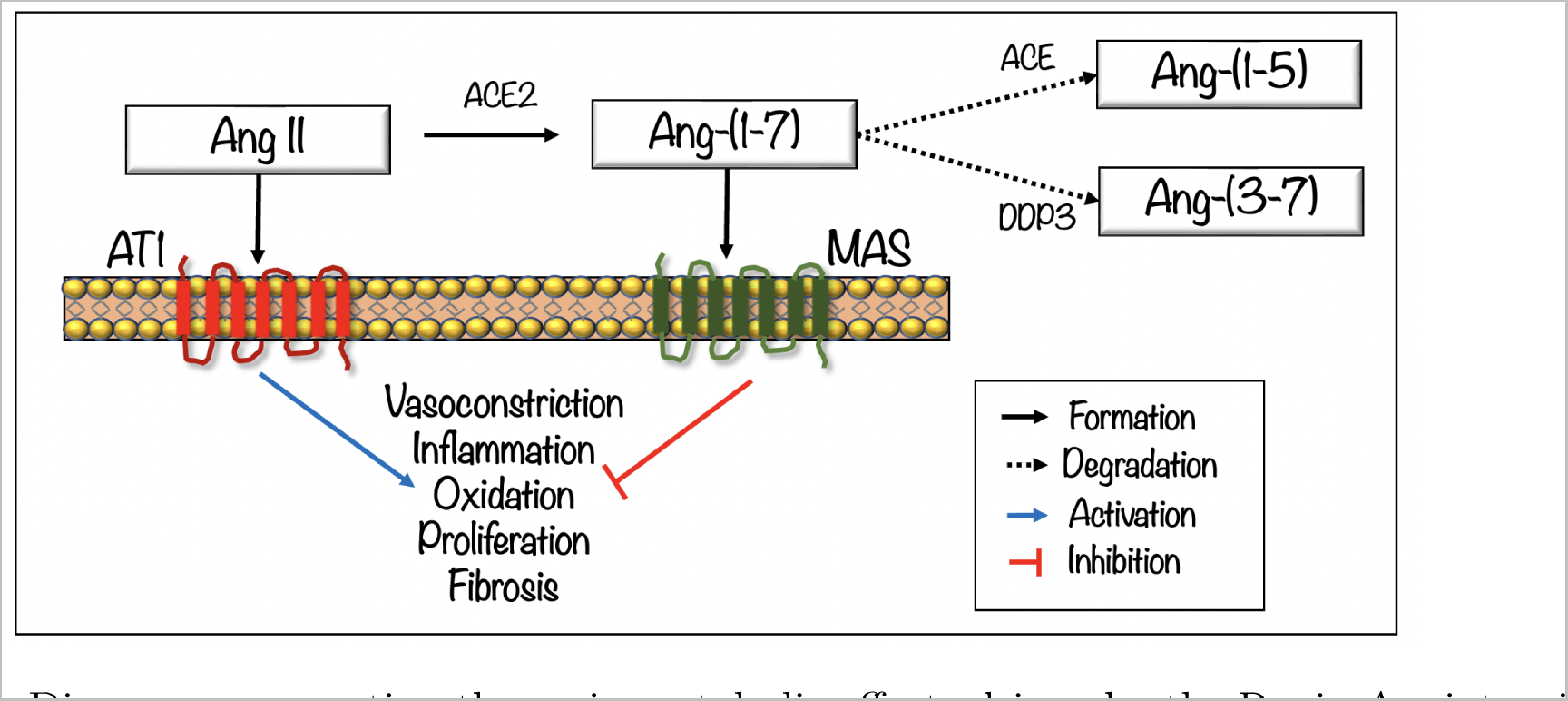
Diagram representing the main metabolic effects driven by the Renin Angiotensin System bioactive peptides (Ang II and Ang-(1-7)) when interacting with its cell receptors AT1 and MAS, respectively. The main proteolytic cleavage of Ang-(1–7) by angiotensin-converting enzyme (ACE) and dipeptidyl peptidase 3 (DPP3), which is associated with Ang-(1-7) rapid degradation and limited therapeutic application.

Recently, Ang-(1-7) has emerged as a novel therapeutic agent who acts as a RAS modulator. It has been proved to be a safe and efficacious (intravenous) IV drug, and since it is not pathogen-specific, it is thought to support a natural healing process.^4^ Several clinical and preclinical studies showed that it can protect organs like lung, kidney, and heart.^4^ Also, some clinical trials testing the safety and the efficacy of intravenous Ang-(1-7) infusion in COVID-19 patients with severe pneumonia are currently under investigation. ^8^ However, Ang-(1–7) has faced limited clinical application due to its unfavorable pharmacokinetic profile: short half-life, poor systemic distribution and poor ability to cross physiological barriers as well as a need for controlled dosage.^9–12^ To overcome these challenges, several strategies have arisen: Ang-(1-7) key aminoacid substitution with a cyclic non-natural amino acid,^11^ cyclization of Ang-(1-7) by inclusion of a thioether bridge,^12, 13^ Ang-(1-7) inclusion into *β*-cyclodextrin molecule,^14^ liposome-entrapped of angiotensin-(1-7)^15^ and an Ang-(1-7) eluting polymer-coated medical device.^16^ One of the most promising approaches is the physically entrapment of a drug inside a dendrimer-based nanocarrier using non-covalent interactions.^17, 18^ Dendrimers are strong candidates because they offer structural stability, high carrying capacity of molecules, water solubility, modifiable surface functionality, available internal cavities, pH dependence properties and a high capacity to cross cell barriers together with an ease of synthesis or commercial availability.^19^ Most of the previous studies have focused on PAMAM, among all dendrimers and hydroxyl-terminated PAMAM dendrimer has been additionally proven to be a non-cytotoxic variant compared with the original amino-terminated PAMAM dendrimer that shows a slight cytotoxic effect.^20^ By using the latter strategy, the capacity of PAMAM-OH dendrimer as an Ang(1-7) carrier was previously explored by means of theoretical and experimental studies^21^ and main conclusions pointed to: a) one dendrimer could be able to capture two peptides; b) Ang-(1-7) could be protected by the dendrimer in a range between 50-65 %; c) Ang-(1-7)/dendrimer forms stable complexes over 60 ns MD which is confirmed by electrophoretic shift assay; and d) Ang-(1–7)/PAMAM-OH complex had an anti-atrophic effect when administered intraperitoneally in mice. We now extend the strategy to design Ang-(1–7)/PAMAM-(OH/NH) complexes by using Ang-(1-7) NMR structure instead of a molecular model, performing molecular docking taking into account the dendrimer and the peptide flexibility, considering different pH conditions in order to glimpse a possible loading and unloading mechanism and extending sampling of the systems.

In summary, we coupled Ang-(1-7) to the most studied kind of dendrimers, PAMAM dendrimers, in order to evaluate their capacity to improve the Ang-(1-7) pharmacokinetic properties and to gain insight into the host-guest (dendrimer/Ang-(1-7)) molecular binding. Understanding the basis of the host-guest molecular binding is fundamental when a very specific and controlled drug release is needed. It is important to note that the beneficial effect of Ang-(1–7) peptide take place only at low concentrations and overdosing might interfere with its receptor-associated functions. ^12^ Different pH conditions (acidic and neutral) and two different group terminals (amino and hydroxyl) were tested as they are critical factors for modulating drug delivery and cytotoxicity. Rigorous computational approaches, such as molecular dynamics (MD) and metadynamics (MTD) have been used to evaluate the structural and energetic properties of the Ang-(1-7), dendrimer or the new complexes formed. Finally, we propose that the best complexes found can be experimentally tested to validate the properties predicted through *In Silico* approaches, as a potential therapeutic agent.

## 2 Methods

### Molecular models

The NMR structure of Ang-(1-7) peptide (ASP-ARG-VAL-TYR-ILEHIS-PRO) in solution was retrieved from the protein database with ID: 2JP8; the topology was created using the pdb2gmx module of GROMACS 5.0.7^22^ and its parameters were downloaded from the GROMOS-compatible 2016H66 force field;^23^ protonation states were assigned according to physical-chemical properties predicted with peptide calculator server^24^ and verified by using ProPkA software^25^ (zero peptide charge for neutral pH and +2 for acidic conditions) as shown in the Figures S1-S3 of the Supplemental Material.

Two types of PAMAM dendrimers were considered: Generation 4 amino-terminated ethylenediamine (EDA)-cored poly(amidoamine) dendrimer (PAMAM-NH_2_) and generation 4 hydroxyl-terminated ethylenediamine (EDA)-cored poly(amidoamine) dendrimer (PAMAM-OH) as depicted in Figures S4-S5. The initial structures and topologies of the dendrimers were created using the newly developed dendrimer topology builder called pyPolyBuilder.^26^ The parameters of the dendrimer and its partial charges were obtained from the GROMOS-compatible 2016H66 force field.^23^ Previously, systematic evaluation of the accuracy of this force field was performed in the simulation of PAMAM dendrimers.^27^ Characteristics of the dendrimers building-blocks used in this work can be found in Table S.5-S.17 of the Supporting Information from the work of Ramos et. al.^27^ and in Table S1 of this work. Two pH values were considered: acidic pH of 3 and neutral pH of 7. To mimic pH conditions, the protonation states for the PAMAM dendrimers were assigned according to the experimental work by Cakara et al.:^28^ at neutral pH, the primary amines were considered protonated and at low pH, both primary and tertiary amines were considered protonated (Figure S6).

### MD of isolated molecules in solution

Dendrimers and peptide structures were placed separately in a box of SPC water model and then subjected to minimization, equilibration and production of standard MD simulations using GROMACS 5.0.7^22^ software, to obtain initial relaxed structures of both types of molecules. Systems were named as: PNH*^n^*, POH*^n^*, PNH*^a^* and POH*^a^*, Ang-(1-7)*^n^* and Ang-(1-7)*^a^*, where letters “n” and “a” correspond to neutral pH and acidic pH, respectively. Abbreviations PNH and POH correspond to amino-terminated PAMAM and hydroxyl-terminated PAMAM dendrimers, respectively.

### Clustering

Representative structures were selected from each MD using clustering by conformations. In the case of dendrimers, the middle structure of the 3 largest clusters was taken as the representative conformation, meanwhile, for peptide, the middle structures in all of the obtained clusters were used unless the cluster had less than 10 members.

### Molecular docking

A double molecular docking was performed in order to have initial dendrimer/peptide complexes by using Autodock Vina^29^ program. We tested all the conformers of the dendrimer and the peptide previously selected. First, a blind rigid docking was performed separately for all combinations of the conformers (each dendrimer conformer is docked to each peptide conformer). Once the peptide conformer found a binding site, a second flexible local docking was performed in this area for each system allowing a better adaptation in the binding site. According to the number of hydrogen bonds (HBs) and Vina affinity, one peptide conformer was selected for each dendrimer site avoiding overlapping. We found that peptides can interact with 3 up to 4 different dendrimer sites for both types of terminal groups. Thus, we decided to choose a configuration with 3 peptides because these peptide conformers fulfill better the above-mentioned criteria, it would be interesting for future works to systematically test the number of peptides attached to each dendrimer. Detailed results are presented in supplemental material (Table S2). The double molecular docking approach was developed using our in-house bash script algorithm.^30^

### MD for complexes

A multi-ligand complex was added to each dendrimer type MD simulation and at each pH value. A total of 4 complexes: [PNH-A]*^n^*, [POH-A]*^n^*, [PNH-A]*^a^* and [POH-A]*^a^*, where letters correspond to neutral pH (n), acidic pH (a) and amino-terminated PAMAM (PNH), hydroxyl-terminated PAMAM (POH) and ang-(1-7) peptides (A). Each complex was placed separately in a SPC model water box and then the three steps of energy minimization, equilibration and production of standard MD were performed using GROMACS 5.0.7 software.^22^

### Metadynamics

In metadynamics (MTD), an external history-dependent bias potential is constructed in the space of a few selected degrees of freedom namely the collective variables (CVs), the effect of the metadynamics bias potential being to push the system away from local minima into exploring new regions of the phase space.^31^ In the long time limit, the bias potential converges to minus the free energy^32^ (or a proportional quantity in the case of well-tempered metadynamics (WT-MTD))^33^ as a function of the CVs. WT-MTD allows an exhaustive sampling of all the possible conformations assumed by the molecules in solution, showing its real flexibility. For Ang-(1-7) metadynamics, we first used the R*_g_* as CV, however, it was not enough to describe the peptide behaviour in solution and led to a stucked system in low R*_g_* values as depicted in Figure S7. In our final set up, we used the radius of gyration (R*_g_*) and the total number of donor-acceptor contacts (*C*_d-a_) as collective variables because they are appropriate descriptors of the behavior of flexible molecules in solution. ^34^ *C*_d-a_ was calculated using a switching functions inspired by the work of Barducci et al,^35^

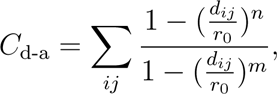

where *r*_0_ was set to 0.25 nm and *n* and *m* were set to 6 and 12, respectively. The sum runs over all of the pairs *i, j* of Hydrogen, Nitrogen and Oxygen polar atoms and *d_ij_* is the distance between each *i* and *j* atoms. In order to limit the R*_g_* sampling, an upper and a lower harmonical restraint was applied. The restraining potential is in the form:

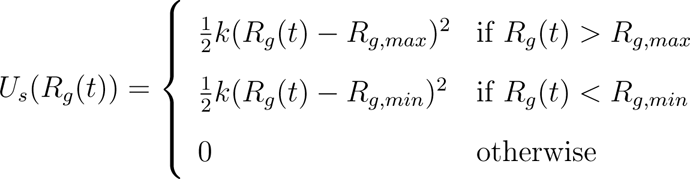

where *k* has to be large enough to prevent the peptide to get stuck in an extended/compressed conformation, *R_g_* (*t*) is the value of the *R_g_* at time *t*. *R_g,max_* and *R_g,min_* are the maximum and minimum values of the *R_g_* that we are interested in.

## 3 Computational details

During MD simulations each system was placed in a cubic simulation box such that all dendrimer/peptide/complex atoms were, at least, 1 nm distant from the box edges. The box was then filled with a sufficiently large number of SPC water molecules. The MD simulation box was neutralized by adding the appropriate number of Cl*^−^* and Na^+^ counterions at a physiological concentration (0.15 M). Systems were subsequently energy minimized under periodic boundary conditions. The equilibration of each system was first performed under the NVT canonical ensemble for 1000 ps. Initial velocities were generated according to a Maxwell-Boltzmann distribution corresponding to a temperature of 300 K. Pressure coupling was then switched on and the system was equilibrated under the NPT isothermal–isobaric ensemble for 1000 ps with a reference pressure of 1 bar. Production runs were carried out under NPT ensemble at 298.15 K and 1 bar for 200 ns. The coordinates were written to the output file every 10 ps for the final analysis. Newtonian equations of motion were integrated using the leapfrog scheme with a time step of 2 fs. Periodic boundary conditions were adopted in all cases. All bond lengths were constrained to their reference values employing the LINCS algorithm.^36^ Electrostatic interactions were calculated with the PME method.^37^ Atomic-based pairlist generation was used as implemented in the Verlet algorithm. When needed V-rescale^38^ thermostat at 298 K and Parrinello-Rahman^39^ barostat at 1 bar were used. This protocol is based on a systematic study that has been recently carried out, confirming that the present scheme is valid.^27^ Details on the molecular systems considered in the present work are presented in Table 1 and Table 2.

**Table 1:** Isolated molecular systems considered in the present work.

| Type | GN | pH | Molecule atoms | Total atoms | Cl<sup>-</sup> | Na<sup>+</sup> | Water molecules |
| --- | --- | --- | --- | --- | --- | --- | --- |
| PNH<sup>n</sup> | 4 | neutral | 1312 | 138360 | 192 | 128 | 45576 |
| PNH<sup>a</sup> | 4 | acid | 1374 | 125148 | 243 | 117 | 41138 |
| POH<sup>n</sup> | 4 | neutral | 1184 | 118853 | 109 | 109 | 38917 |
| POH<sup>a</sup> | 4 | acid | 1246 | 122457 | 176 | 114 | 40307 |
| Ang-(1-7)<sup>n</sup> | - | neutral | 85 | 5105 | 5 | 5 | 1670 |
| Ang-(1-7)<sup>a</sup> | - | acid | 87 | 5136 | 7 | 5 | 1679 |

**Table 2:**
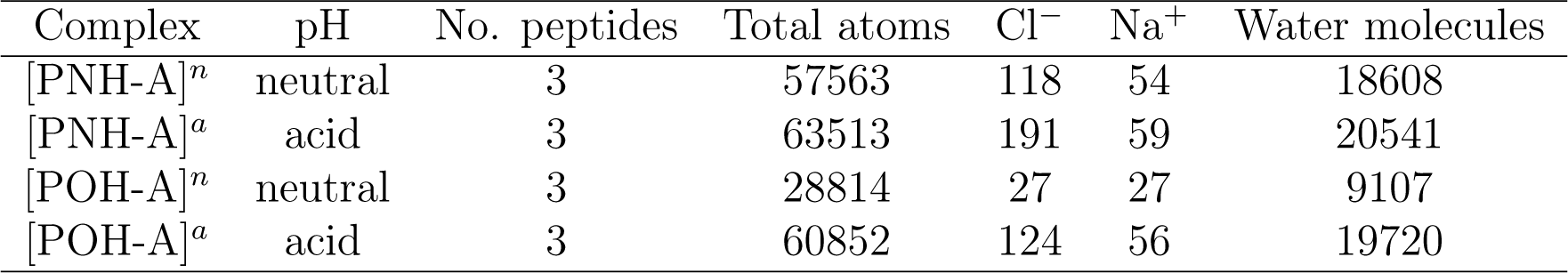
Complexed molecular systems considered in the present work. P in the name corresponds to PAMAM, NH for amino terminated dendrimer, OH for hydroxyl terminated dendrimer, A for Ang-(1-7) followed by the number of peptides in complex with the dendrimer and the last number that identify the two complex conformers for each system.

During clustering by conformation, we used the Gromos clustering algorithm by Daura et al.^40^ During docking, the binding mode that shows the lowest energy was chosen. The grid box during rigid docking was centered on the dendrimer center with a cubic grid box of side 26 Å. The grid box during flexible docking was centered on the site that found in the first docking with a cubic grid box of side 10 Å.

In Ang-(1-7) MTD, we used gaussian width of 0.1 nm, gaussian height of 1.2 kJ/mol, deposition time between gaussians of 500 MD time steps, a biasfactor of 15 and a temperature of 300 K. Convergence was reached in 1 *µ*s according to the behaviour of c(t) (a time-dependent parameter in the definition of the free energy which makes the free energy asymptotically time-independent) and the diffusion of the collective variable^41^ as presented on Figure S7. Gromacs-2016.3 and Plumed-2.3.3 were used.

## 4 Results and Discussion

### 4.1 MD of isolated systems in solution

#### 4.1.1 PAMAM dendrimers

The Root-mean-square deviation (RMSD) of atomic positions is a standard measure of the structural changes mainly used for proteins or non-protein small organic/inorganic molecules. Nevertheless, due to the large conformational flexibility of dendrimers in aqueous solution, there is a debate as to whether it is an appropriate molecular descriptor to account for its structural stability. Here, we have employed RMSD analysis to test the convergence of the simulations, in order to obtain equilibrated initial structures for the following steps. As shown in Fig 2a, the PNH*^n^* RMSD values undergo large changes from the starting structure in the first 50 ns, afterwards the system seems to reach a stable behavior; meanwhile PNH*^a^* RMSD values undergo big changes from the starting structure in the first 20 ns, afterwards the system seems to reach a stable behavior fluctuating around a mean value. The POH*^n^* RMSD values undergo a rapid change during the first ns, after that, the RMSD steadily deviates slowly from this new conformation, reaching a new steady-state after 100 ns; meanwhile, POH*^a^* RMSD values undergo big changes from the starting structure in the first 20 ns, afterwards the system seems to reach a stable behavior fluctuating around a mean value. Ang-(1-7)*^n^* and Ang-(1-7)*^a^* RMSD values show a more stable behavior from the first ns. According to these results, we agreed to use the last 100 ns of the simulations as equilibrated data for all the systems. Large changes observed on dendrimer structures are mainly due to the fact that the initial structures were obtained by PyPolyBuilder algorithm which only gives us an initial guess for the dendrimer geometry in vacuum.

**Figure 2:**
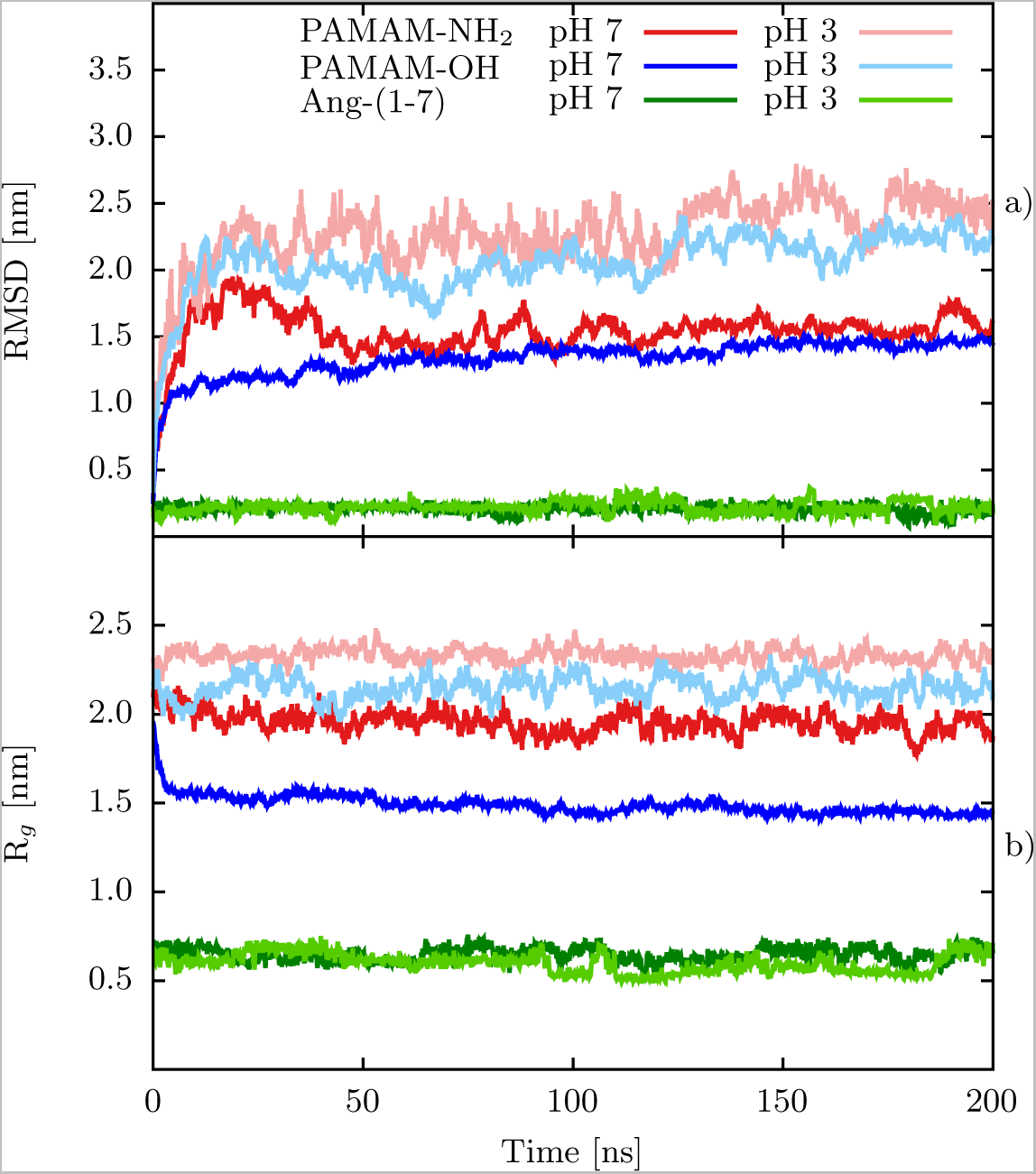
Structural features of PAMAM-NH_2_, PAMAM-OH and Ang-(1-7) systems. a) RMSD and b) R*_g_* values as function of time.

The radius of gyration (R*_g_*) might be a more appropriate molecular descriptor to account for the structural stability of dendrimers, it gives a measure for the compactness of a molecular structure. As shown in Figure 2b, the PNH*^n^* and PNH*^a^* R*_g_* values show a more stable behavior from the first nanoseconds around a mean value. POH*^n^* and POH*^a^* R*_g_* values show that the dendrimer undergoes a rapid change during the first ns around a mean value; finally, Ang-(1-7)*^n^* and Ang-(1-7)*^a^* R*_g_* values show a more stable behavior from the first nanoseconds around a mean value in both cases.

PNH*^n^* and PNH*^a^* R*_g_* mean values in the equilibrated zone were 1.94 *±* 0.05 nm and 2.3 *±* 0.04 nm, respectively. On the other hand, POH*^n^* and PNOH*^a^* R*_g_* mean values in an equilibrated zone were 1.46 *±* 0.02 nm and 2.16 *±* 0.06 nm, respectively. Results show that dendrimers at acidic pH assume a stretched, open configuration similar to the dense-shell model as a consequence of the strong Coulomb repulsion between the charged tertiary/primary amines present in the structure. R*_g_* values are in very well agreement with experimental and theoretical results available,^27, 42–47^ as also compared in Table 3, confirming that GROMOS-compatible 2016H66 force field, together with our chosen methodology were capable to represent the structural properties of the dendrimers used in this work. It can be observed that POH*^n^* and POH*^a^* has in general a more compact structure (lower R*_g_*) compared with PNH*^n^* and PNH*^a^*, respectively (Figure 2b). At neutral pH, this is partly due to the preference of the group terminals to be surrounded by waters (PNH*^n^*, Figure 3a) or to interact with internal dendrimer groups (POH*^n^*, Figure 3b), from the total possible interactions between group terminals and waters/internal dendrimer groups, we found that PNH*^n^* forms 95 % of the interactions with waters and 5 % with internal groups, meanwhile, POH*^n^* forms 85 % of interactions with waters and 15 % with the internal groups. The R*_g_* and RMSD fluctuations in the case of PNH*^n^*, PNH*^a^* and POH*^a^* throughout the simulation time, compared to that of POH*^n^*, suggests that the formers have a more flexible structure than the latter. This result might be attributed to the continuous movement of the protonated terminal amine groups in PNH*^n^* and terminal/internal protonated groups in PNH*^a^* and POH*^a^* to minimize the unfavorable enthalpic penalty due to electrostatic repulsion on charges of the same sign.

**Figure 3:**
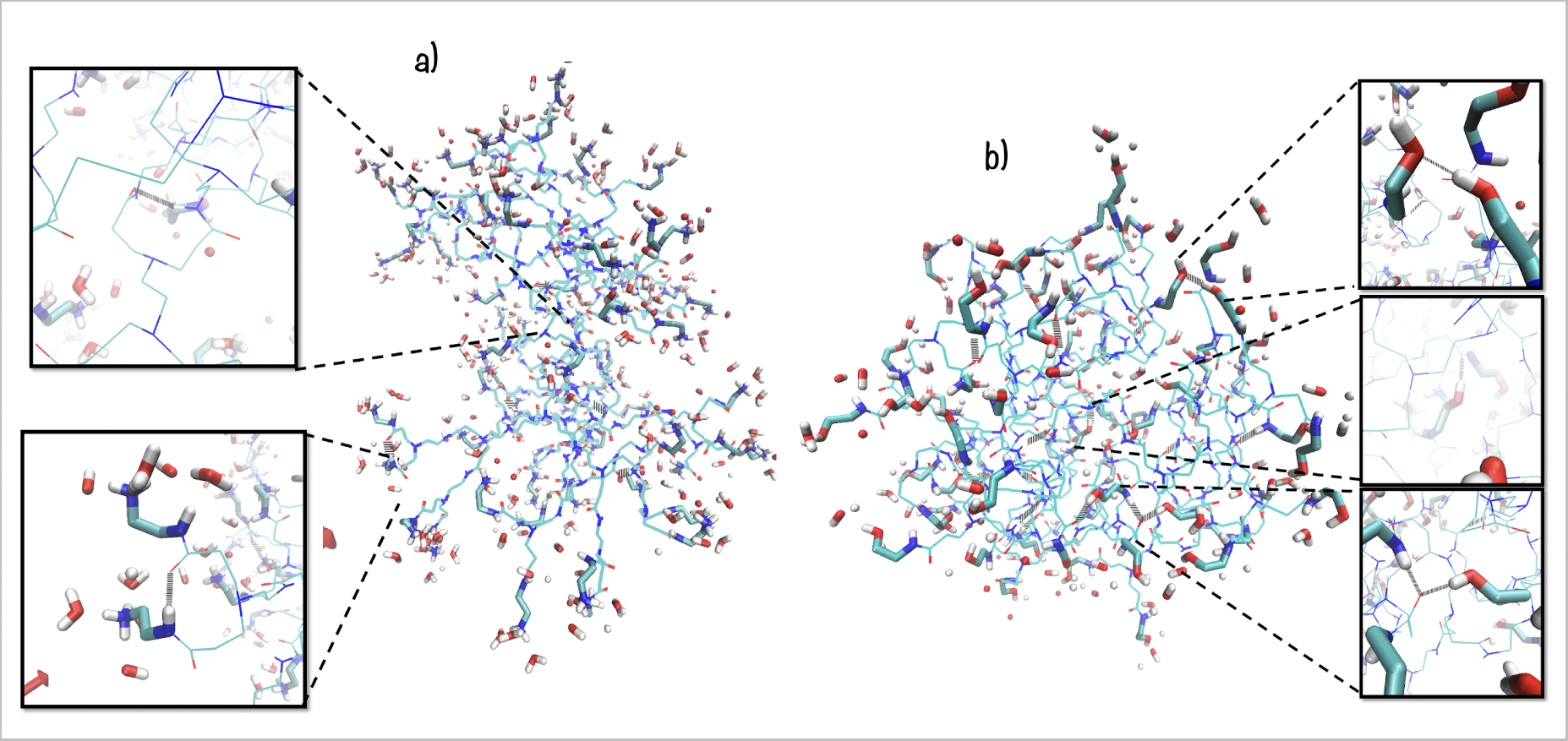
Waters sorrounding group terminals and hydrogen bonds formed between terminal groups and waters/dendrimer internal groups. a) PAMAM-NH_2_ and b) PAMAM-OH, at neutral pH.

**Table 3:**
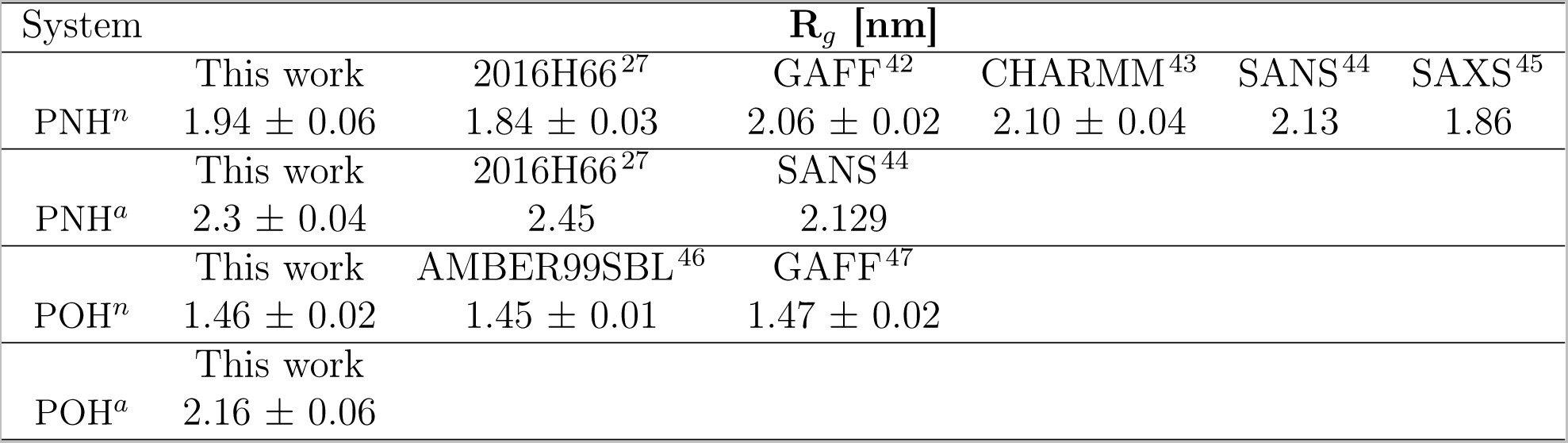
Radius of gyration R*_g_* of PAMAM dendrimers and comparison with experimental and theoretical studies.

| System | $R_g$ [nm] | | | | | |
| --- | --- | --- | --- | --- | --- | --- |
|  | This work | 2016H66<sup>27</sup> | GAFF<sup>42</sup> | CHARMM<sup>43</sup> | SANS<sup>44</sup> | SAXS<sup>45</sup> |
| PNH<sup>n</sup> | $1.94 \pm 0.06$ | $1.84 \pm 0.03$ | $2.06 \pm 0.02$ | $2.10 \pm 0.04$ | 2.13 | 1.86 |
|  | This work | 2016H66<sup>27</sup> | SANS<sup>44</sup> |  |  |  |
| PNH<sup>a</sup> | $2.3 \pm 0.04$ | 2.45 | 2.129 | | | |
|  | This work | AMBER99SBL<sup>46</sup> | GAFF<sup>47</sup> |  |  |  |
| POH<sup>n</sup> | $1.46 \pm 0.02$ | $1.45 \pm 0.01$ | $1.47 \pm 0.02$ | | | |
|  | This work |  |  |  |  |  |
| POH<sup>a</sup> | $2.16 \pm 0.06$ | | | | | |

To characterize the shape of the dendrimers, an asphericity parameter *δ* was estimated. It is important to note that the closer this value is to zero, the more spherical the dendrimer becomes and its calculation is described in detail elsewhere.^27^ As shown in Table 4 and Figure 4, PNH*^n^* assume a less spherical structure compared to POH*^n^*, meanwhile, PNH*^a^* and POH*^a^* show an almost spherical behavior. This high asphericity observed in PNH*^n^*, might be due to an asymmetry induced by a high degree of charged terminal groups back folding as also reported in the previous works,^27^ in contrast, the neutral terminal in POH*^n^* seems to be able to accommodate in a more compact way due to its less charge-charge terminal repulsion and its preference of interactions with internal dendrimer groups. In the case of PNH*^a^* and POH*^a^*, an almost spherical shape might be mainly due to the repulsion between charged tertiary amine groups in the interior of the dendrimers.

**Figure 4:**
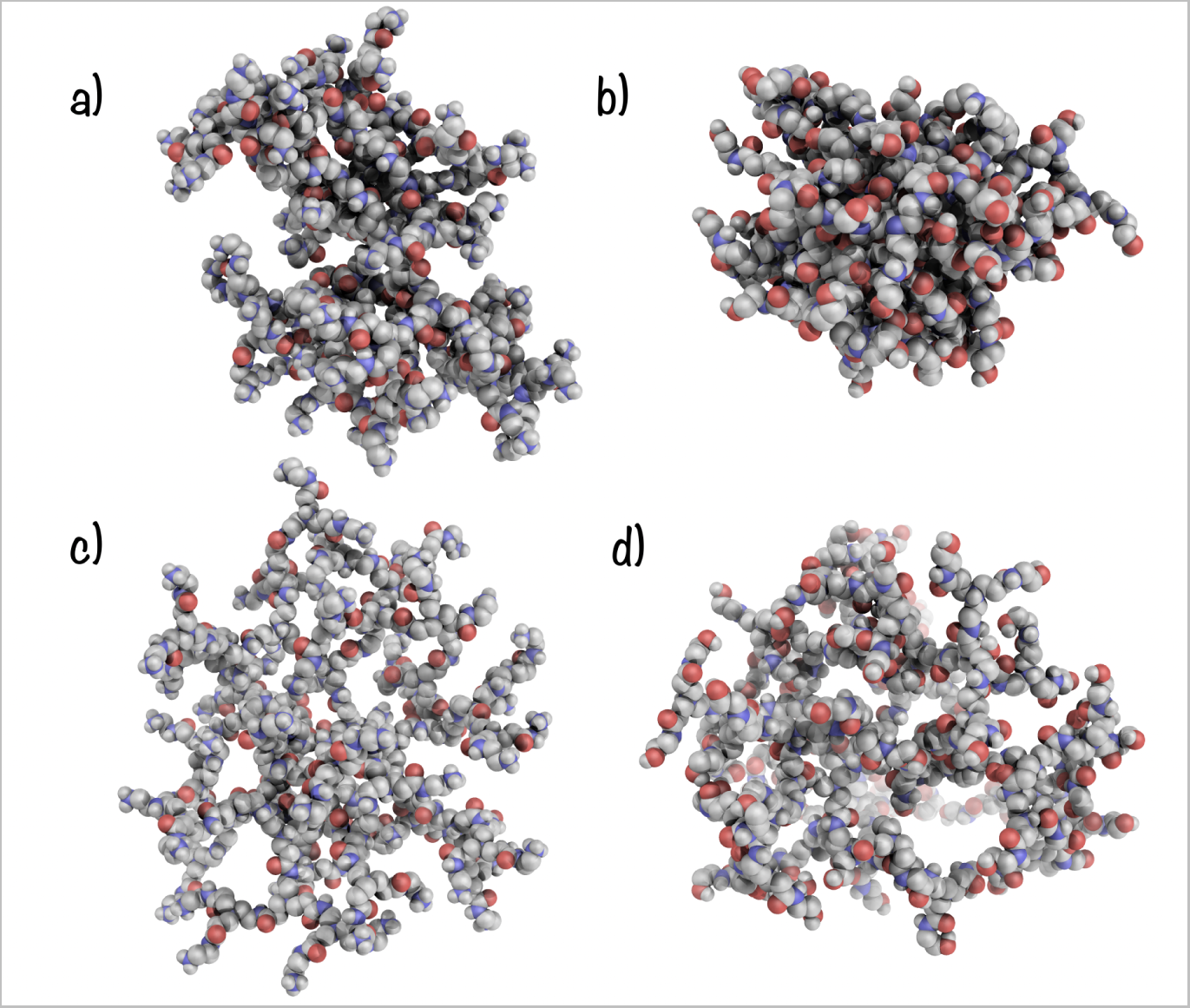
Dendrimers shape comparision. a) PAMAM-NH*^n^*, b) PAMAM-OH*^n^*, c) PAMAM-NH*^a^* and d) PAMAM-OH*^a^* at 200 ns of MD simulation. Leters “a” and “n” states for neutral and acidic pH.

**Table 4:** Asphericity *δ* of PAMAM dendrimer and comparison with theoretical studies available. Leters “a” and “n” states for neutral and acidic pH.

| System | $\delta$ | | | |
| --- | --- | --- | --- | --- |
|  | This work | 2016H66<sup>27</sup> | AMBER<sup>48</sup> | GAFF<sup>42</sup> |
| PNH<sup>n</sup> | 0.04 | $0.08 \pm 0.03$ | 0.02 | 0.03 |
| PNH<sup>a</sup> | 0.004 | $0.02 \pm 0.01$ | 0.01 | |
|  | This work | GAFF<sup>47</sup> |  |  |
| POH<sup>n</sup> | 0.01 | 0.01 |  |  |
| POH<sup>a</sup> | 0.006 |  |  |  |

To characterize how particle density (waters, terminal nitrogens, internal nitrogens and ions) varies as a function of the distance from the dendrimer center, radial distribution functions were measured. As shown in Figure 6a, a higher level of structuration is evidenced by the pronounced peaks on the distributions on PNH*^n^*, PNH*^a^* and POH*^a^* internal nitrogens compared with those of POH*^n^*, this is mainly due to a more dense POH*^n^* dendrimer compared with less dense, PNH*^n^*, being more evident on PNH*^a^* and POH*^a^*, as evidenced by the height of the peaks. As it can be seen, the dense-core model^49^ is confirmed at neutral pH, this model states that the monomer (internal Nitrogen atoms) density distribution is high close to their centers and decays as it approaches the periphery, as evidenced of the height of the peaks from the center to the periphery at neutral pH. However, at acidic pH, dendrimers assume a less dense inner cavity, stretched, open conformation like to the dense-shell model^49^ as a consequence of the strong Coulomb repulsions between the charged units. This is in agreement with theoretical models that predicts that the PAMAM-NH_2_ core becomes denser upon decreasing the acidity of the medium. Interestingly, this could led to a different drug interaction behaviour and load capacity, probably allowing better drug encapsulation in the big inner cavities at acidic pH (Figure 5).

**Figure 5:**
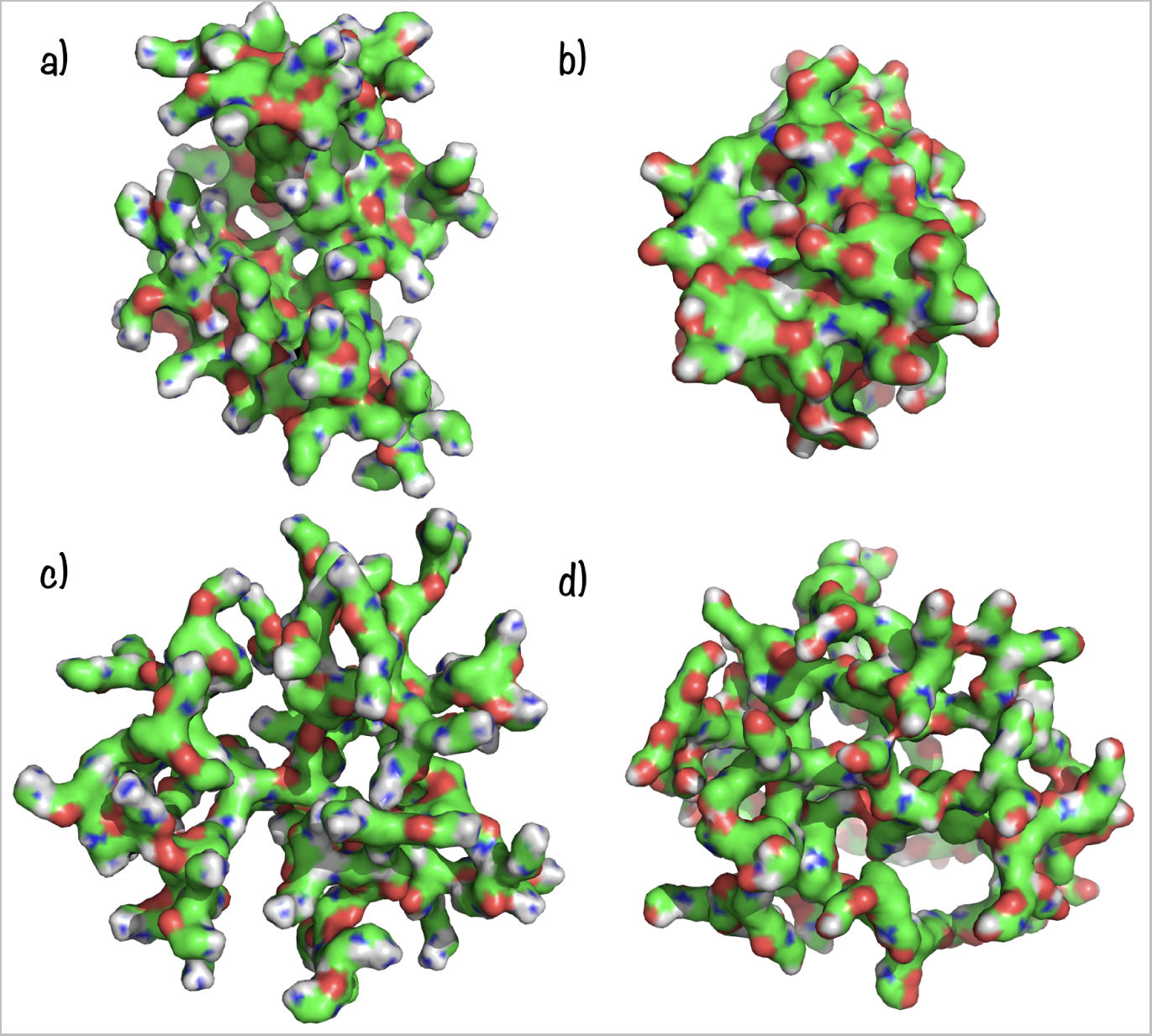
Dendrimers cavities comparision. a) PAMAM-NH*^n^*, b) PAMAM-OH*^n^*, c) PAMAM-NH*^a^* and d) PAMAM-OH*^a^* at 200 ns. Leters “a” and “n” states for neutral and acidic pH.

**Figure 6:**
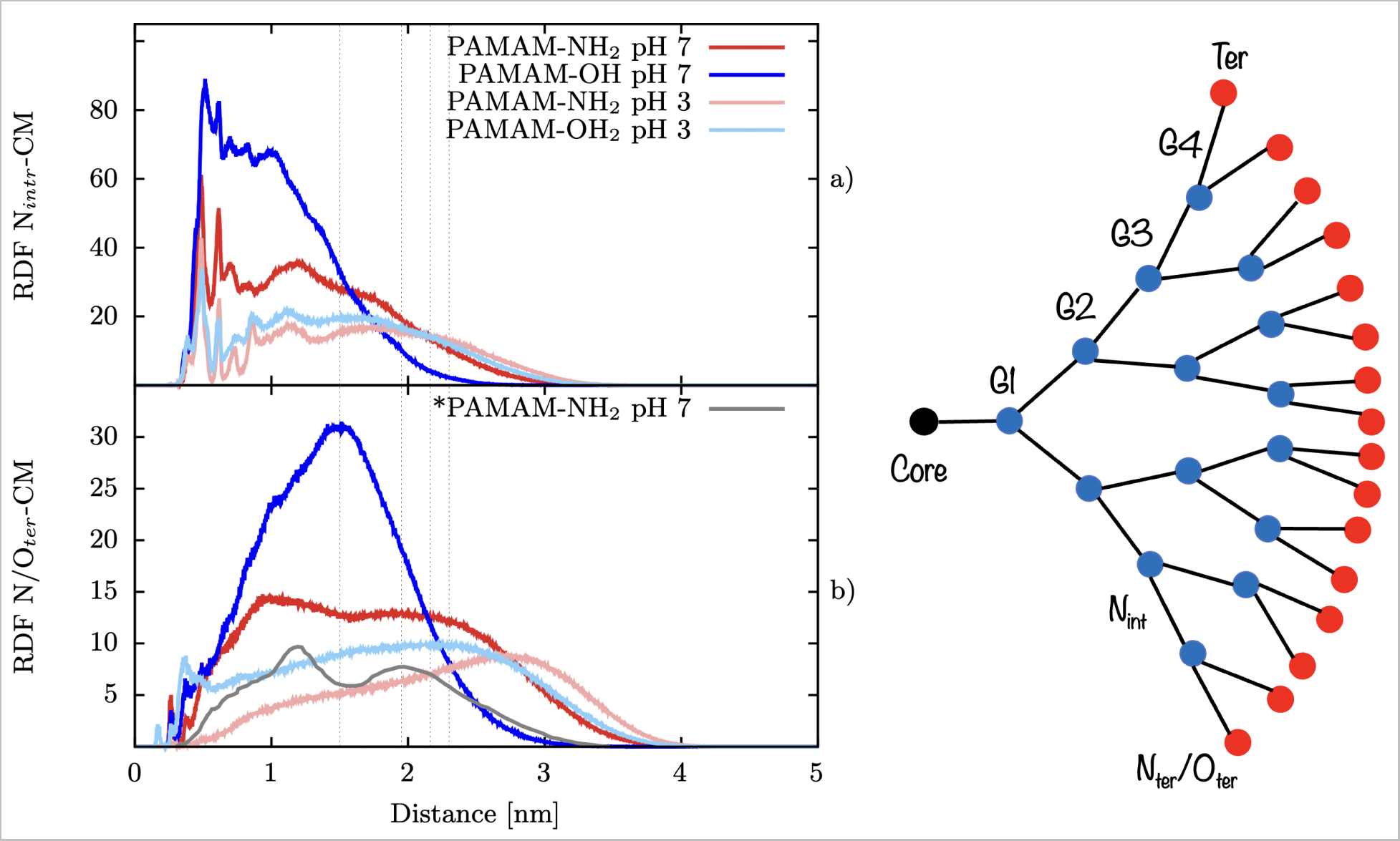
Radial distribution function of molecular groups with respect of the dendrimer COM. a) internal nitrogen atoms and b) terminal nitrogen/oxygen atoms. Schematic diagram at the right of the panel showing the internal and terminal nitrogens. GX states for the dendrimer generation. *Thinner grey line states for comparison from previous work in the literature.^27^ Grey vertical dashed line represents the Rg of the dendrimers.

As shown in Fig 6b, PNH*^n^* N-ter can be found in the intermediate zone (around 1 nm) as well as in the external surface (around 2 nm), compared with the case of PNH*^a^* where distribution is near the external surface (around 2.3 nm), and POH*^n^* and POH*^a^* O-ter which are mainly found on the surface (around 1.5 nm and 2.16 nm). In PNH*^n^*, terminal groups exhibit a broader distribution that might indicate a certain degree of backfolding, being the backfolding increased from low to neutral pH, as shown by the intensity of the density peaks.

As shown in Fig 7a, RDF profiles reveal a major presence of buried water molecules within the structure of PNH*^n^*, PNH*^a^* and POH*^a^* compared with POH*^n^*, at distances larger than 5 Å from the center of mass of the dendrimer. Hydroxyl terminals when internal tertiary amines are not protonated reduce the penetration of water into the dendrimer structure, which can be interpreted as an enhancement of the hydrophobic character of the dendrimer inner small cavities, ideal for encapsulation of small hydrophobic drugs. In general, dendrimers with both kind of terminal groups at acidic pH show a major presence of buried waters whithin its structure in order to solvate the charged groups. As shown in Fig 7b-c, RDF profiles reveal that chloride ions penetrate the innermost dendrimer cavities in PNH*^n^*, PNH*^a^* and POH*^a^*, contrary to the case of POH*^n^*. Interestingly, a broad peak around 1 nm and a shoulder around 2 nm, where PNH*^n^* positive terminals are localized, was found. Sodium counterions are found as far as posible of charged terminals in PNH*^n^*, PNH*^a^* and POH*^a^* and more disperse in POH*^n^* around its surface.

**Figure 7:**
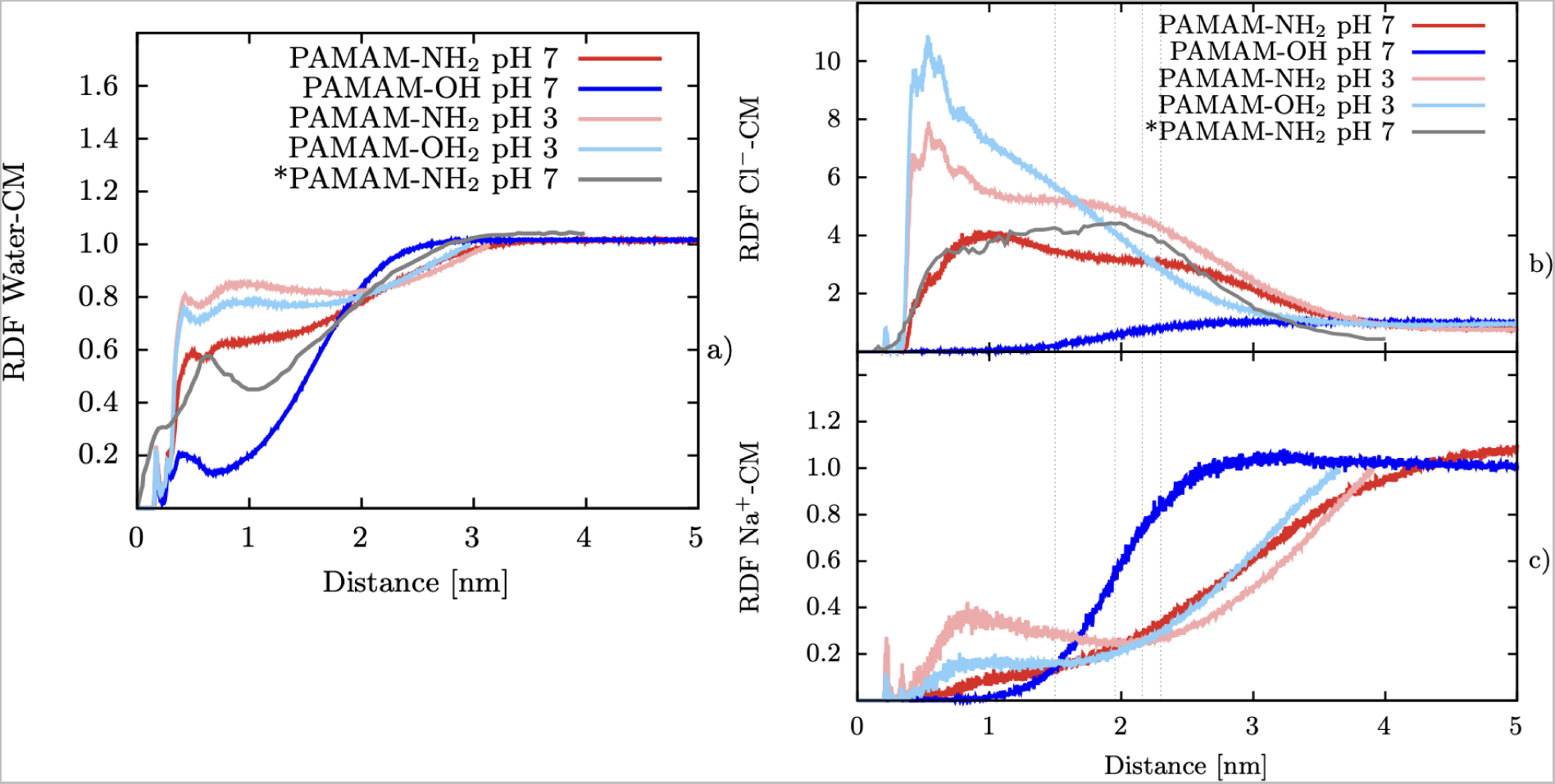
Radial distribution function of molecules with respect of the dendrimer COM. a) water molecules and b) Cl*^−^* ions and c) Na^+^ ions. *Thinner grey line states for comparison from previous work in the literature.^27^ Grey vertical dashed line represents the Rg of the dendrimers.

#### 4.1.2 Ang-(1-7) bioactive peptide

Ang-(1-7)*^n^* and Ang-(1-7)*^a^*, R*_g_* mean values in the equilibrated zone were 0.65 *±* 0.04 nm and 0.57 *±* 0.05 nm, respectively (Figure 2b). From the structural point of view, the solution structure of Ang-(1-7) was previously determined^50^ by means of NMR and circular dichroism experiments performed at acidic pH. There, the authors claimed that Ang-(1–7) structure showed a conformational equilibrium between the random coil and *β*-sheet (or a mixture of bend structures), with a bend stabilized by interactions between residues VAL3 and TYR4, this is in very well agreement with our results in Figure 8, where we can see that the most frequent secondary motif is bent between VAL3 and TYR4. In our classical MD simulations at acidic pH, we found that 60 % of the structures were similar to the NMR solution structure (RMSD *<* 2 Å when superimposed), meanwhile, at neutral pH, this happened only in 40 % of the frames, thus, this conformation was less frequent at neutral pH (Figure 9).

**Figure 8:**
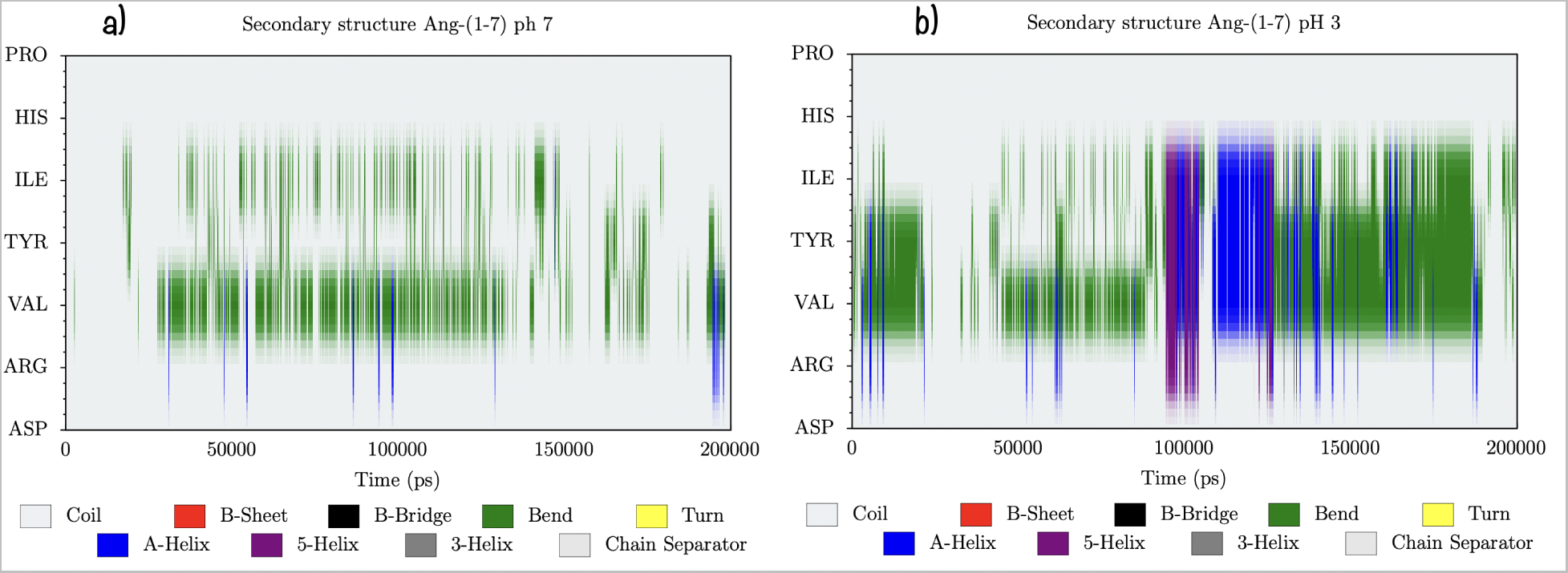
Ang-(1-7) secondary structure as a function of time. a) at neutral pH and b) at acidic pH.

**Figure 9:**
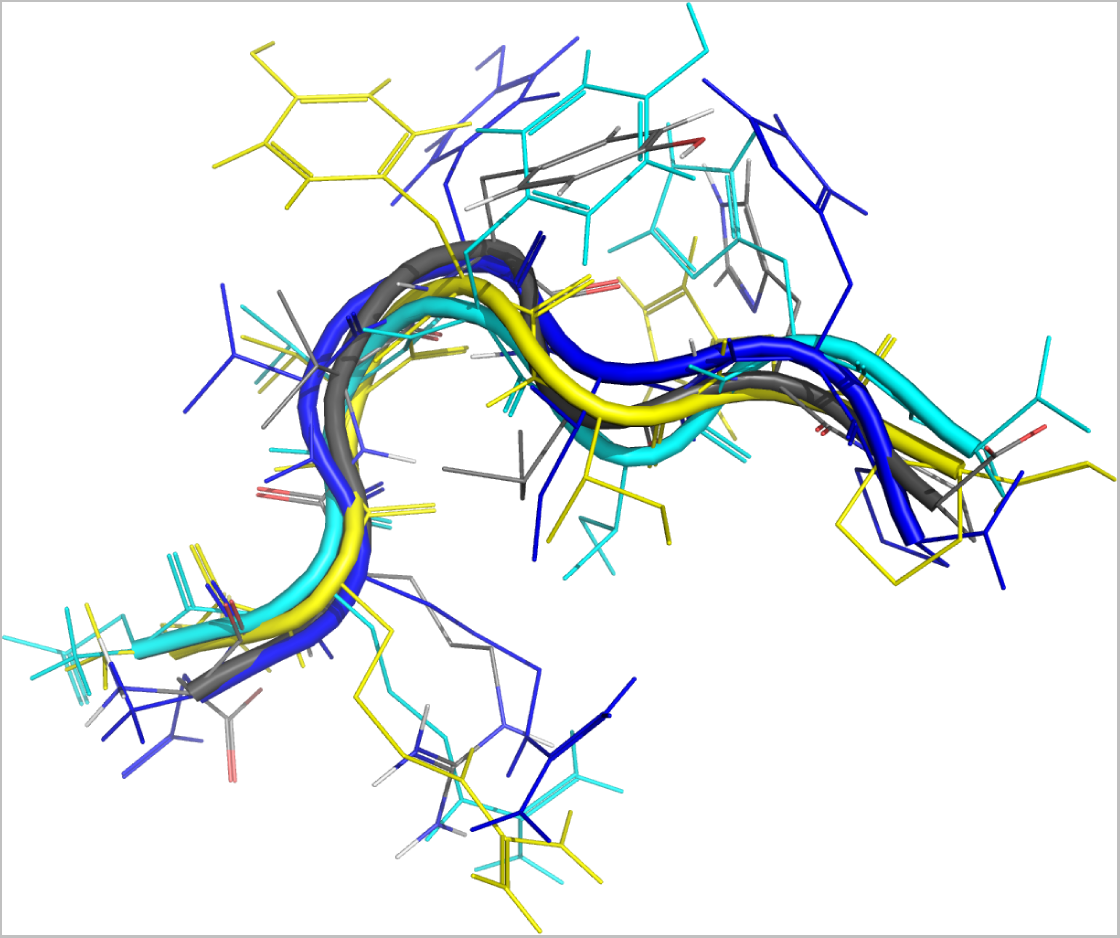
Superimposition of: experimental NMR structure obtained at acidic pH (grey), structure repeated 40 % of the frames in classical MD simulation performed under neutral pH (blue), structure repeated 35 % of the frames in metadynamics simulation performed under neutral pH (cyan), structure repeated 60 % of the frames in classical MD simulation performed under acidic pH (yellow).

The conformational sampling of peptides in solution that are fully or partially disordered and flexible is not always guaranteed with conventional MD due to the limiltations in the sampling time scale.^51^ To get more insights about Ang-(1-7) heptapetide structure and extend this insight at neutral pH conditions, we performed a MTD simulation at neutral pH employing the R*_g_* and the C*_d−a_* as CVs. The final free energy landscape is shown in Figure 10a, where it can be shown that the landscape has a broad and unique minimum or basin A, indicating that this peptide exist in a variety of conformations in solution. The global minima corresponds to a R*_g_* of 0.65 nm and a C*_d−a_* of 30, in very well agreement with R*_g_* values from classical MD; however, the basin A samples a wide range of R*_g_* and C*_d−a_* values, this is a R*_g_* in the region spanning from 0.6 to 0.7 nm and contacts ranging between 30 and 35. Interestingly, the structures in the basin showed a persistent HB between ASP1 and TYR4 or between ASP1 and VAL3 (Figure 10b). This HB was also frequently observed on the bigger cluster of the MD of Ang-(1-7)*^n^* in solution, this conformation is probably frequent because it allows positive charged ARG2 to be far from positive charged N-terminal, both of them becoming able to interact with polar waters.

**Figure 10:**
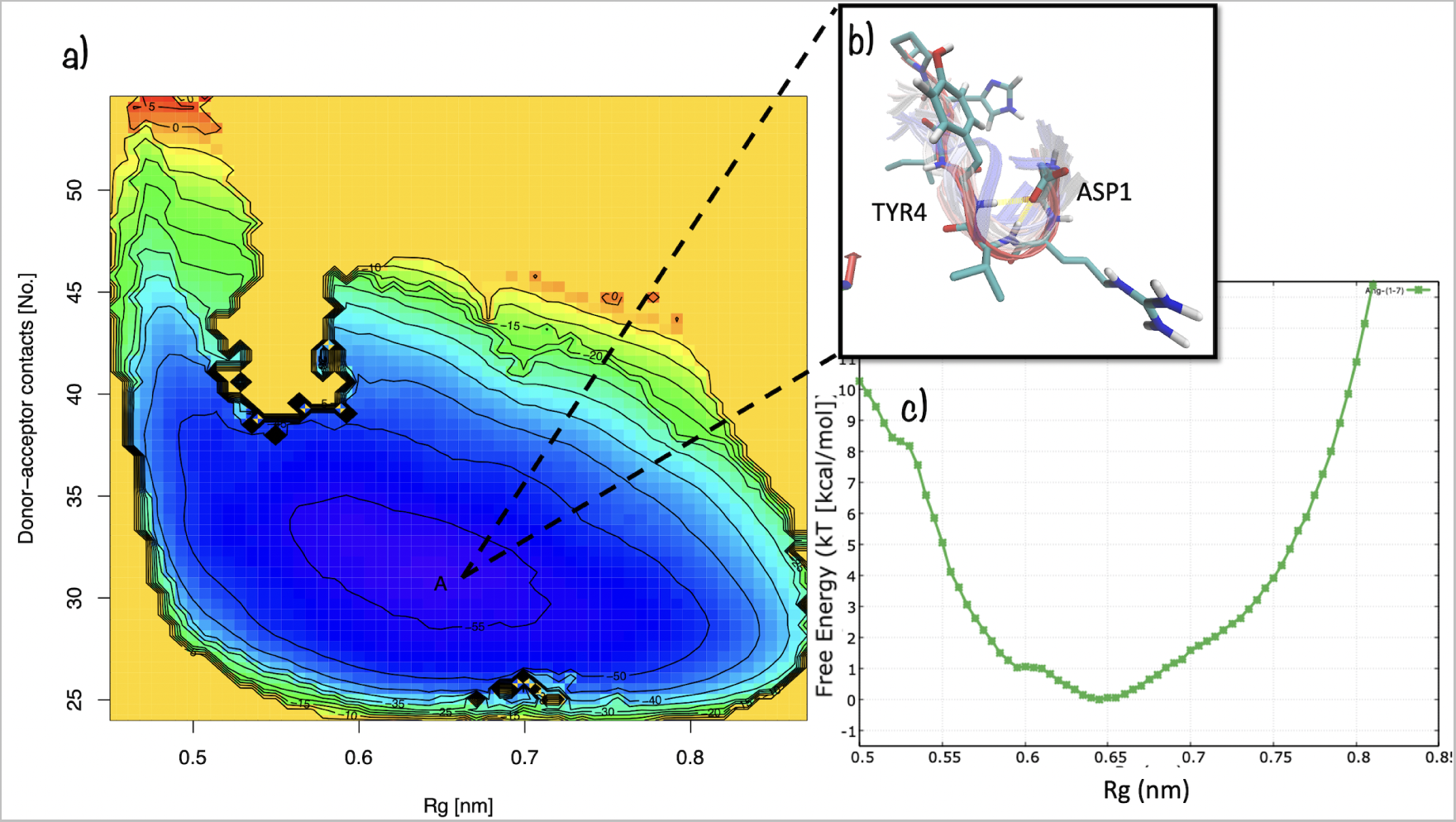
Ang(1-7) 2D a) Free Energy Surface (FES) as a function of the total number of donor-acceptor contacts and the R*_g_* . Isoenergy lines are drawn every 5 kcal/mol. T = 300 K, pH = 7. b) Structures representative of the main free-energy basin A are superimposed in the inset figure. c) Estimate of the free energy as a function of the R*_g_* from a well-tempered metadynamics simulation.

The one dimensional FES as a funcion of R*_g_* is shown in Figure 10c, as can be seen, there are not large free energy barriers (around 1 kT) dividing the metastable states in the main basin A. It is important to note that only 35 % of the structures in basin A were similar to the NMR structure and in agreement with classical MD. This result is coherent with the conformational equilibrium between the random coil and a mixture of bend structures reported experimentally for Ang-(1-7). However, basin A is characterized by a significant degree of flexibility of the N-terminal and C-terminal tails which samples multiple conformations that deviate from the ensemble of the NMR structure as shown in Figure 11.

**Figure 11:**
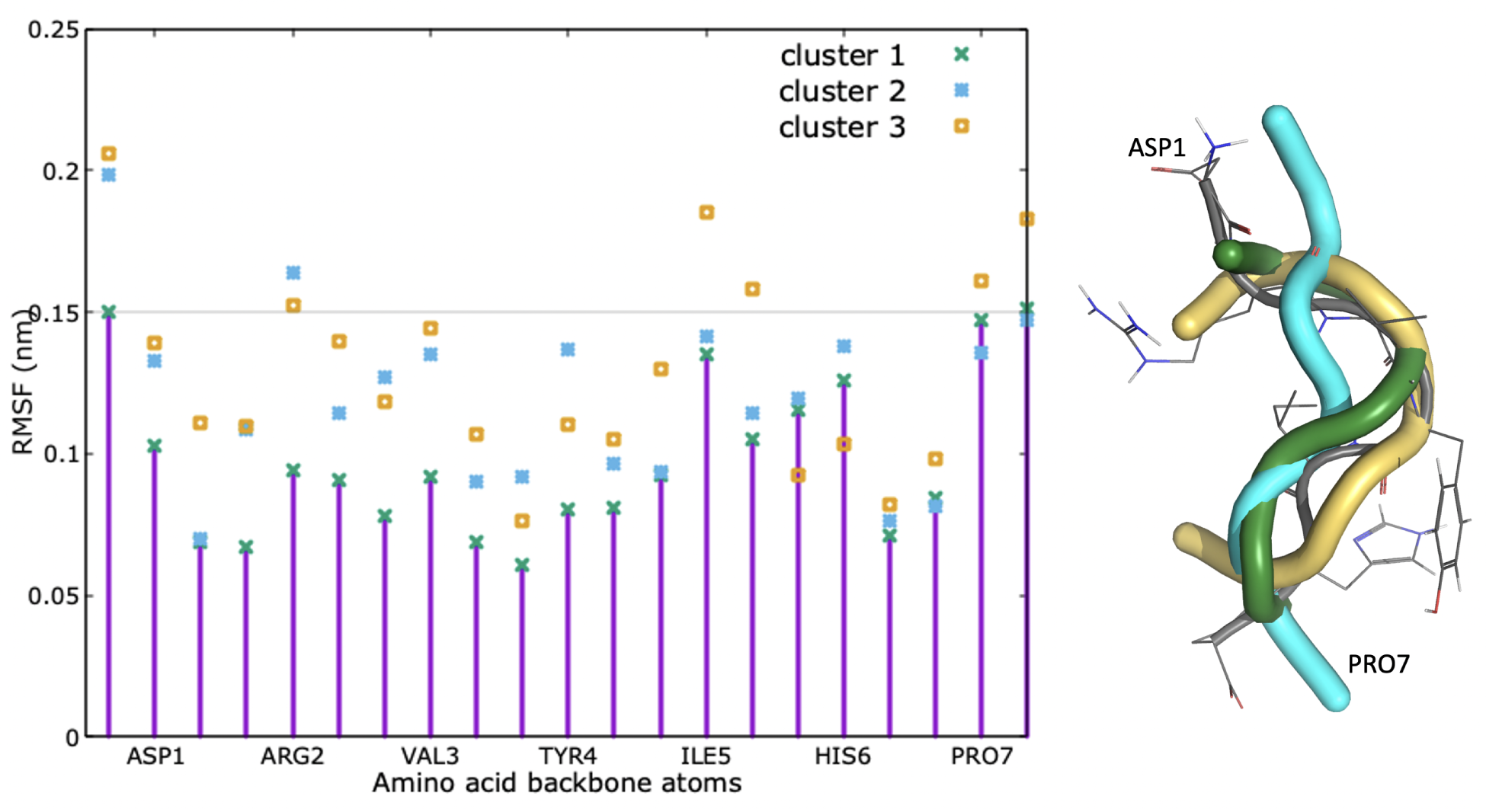
RMSF of Basin A main clusters as a function of aminoacid backbone atoms. Structures representative of the basin A main clusters are superimposed in the right figure. Grey represent the NMR structure. Threshold of 0.15 nm is marked to visualize the most flexible atoms. Cluster 1 is marked also with impulses to guide visualization.

Together, the R*_g_*, *δ* and RDFs results validates our chosen methodology and confirms that GROMOS-compatible 2016H66 force field is capable to model the theoretical and experimental structural features of G4-PAMAM dendrimers with amino or hydroxyl group terminals. In case of Ang-(1-7), structural super impositions of MD and metadynamics structures with NMR structure confirms the experimental results previously reported.

### 4.2 Clusters and docking

The middle structure of the most populated clusters for dendrimers and the peptide were used for the molecular docking of each dendrimer conformer with each of the peptide conformers. Our methodology allowed us to find a suitable binding site for the peptide by using a representative peptide conformation taken from the MD, once the peptide finds a reasonable site, the algorithm allows it to better accommodate into this local site, as can be schematized in Figure 12.

**Figure 12:**
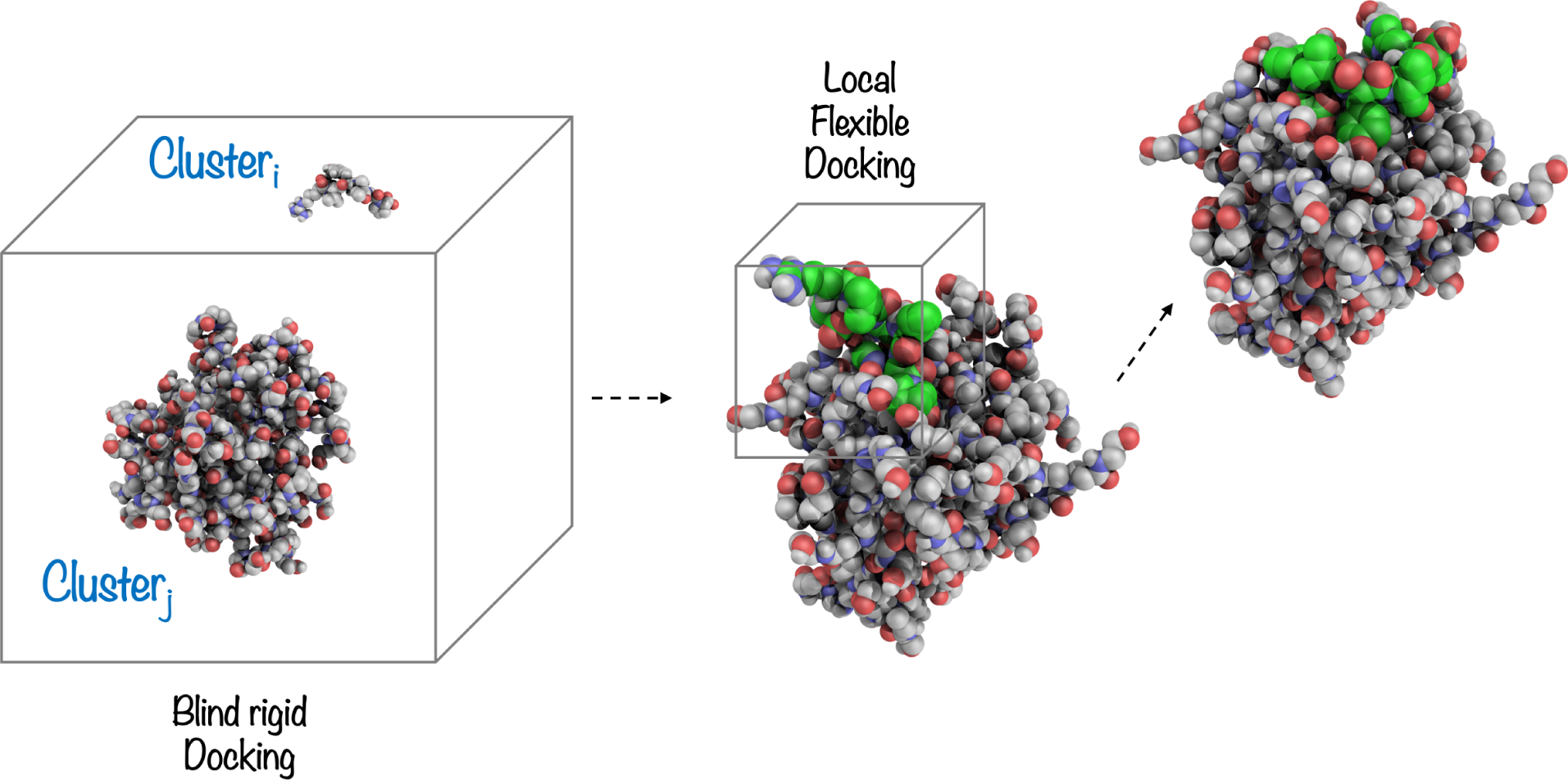
Double-docking schematic representation. Each of the peptides conformer is allowed to find a site by using a blind rigid docking, afterwards, once the peptide found a reasonable site, a flexible docking allows it to better accommodate into the site.

Final complexes resulted from the docking are shown in Figure 13. As can be seen, PNH*^n^* allows the interaction of the peptides with its inner shells as well as with its surface. POH*^n^*, only allow the peptide interactions with its surface due to its more compact structure. We can observe that at acidic pH, dendrimers allow a better encapsulation of 3 peptides. Details on the HBs formed between peptides and dendrimers are presented in Tables 5 and 7.

**Figure 13:**
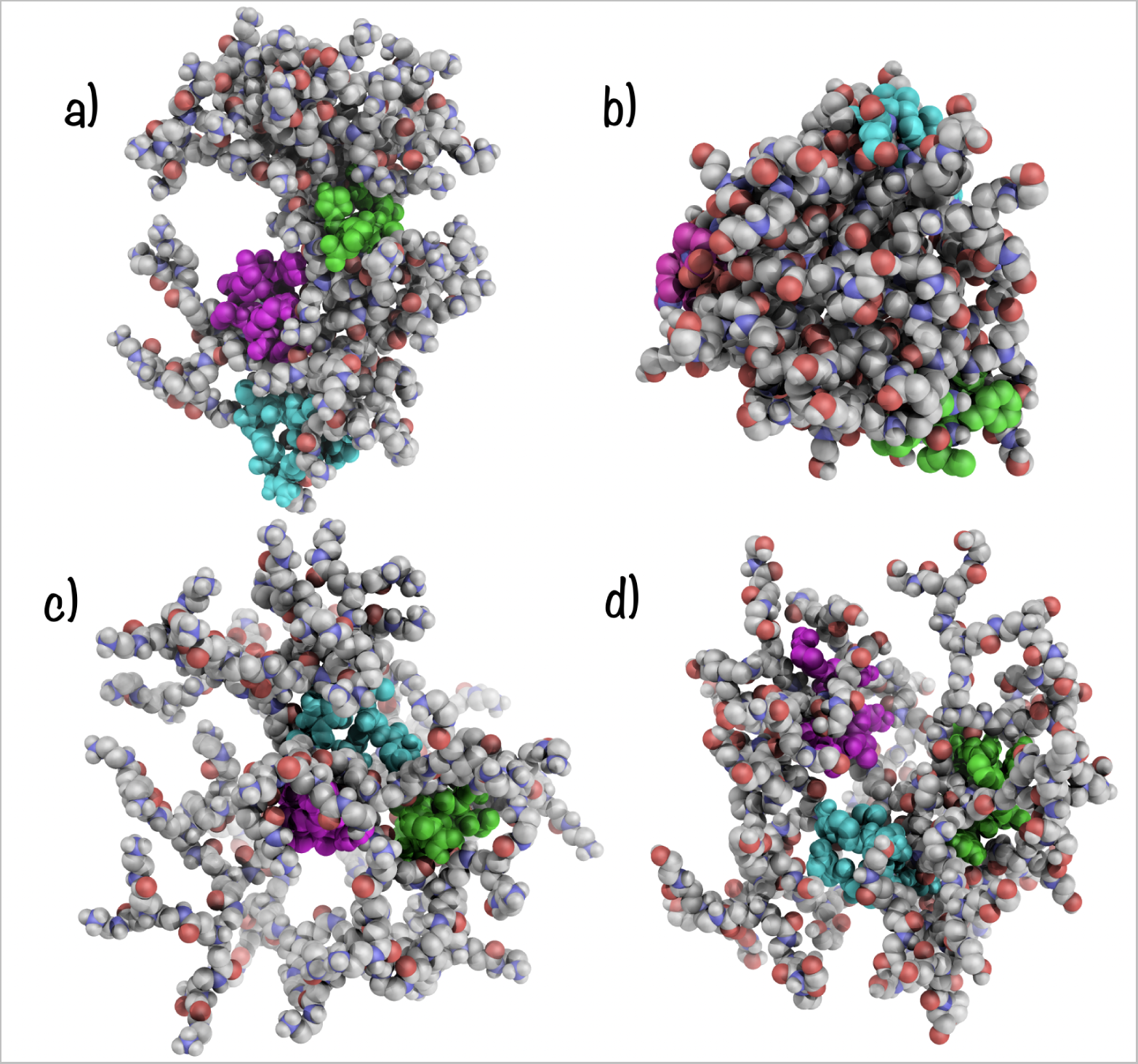
MD complexes initial structures obtained from double-docking approach: a) PNH*^n^* in complex with 3 Ang-(1-7)*^n^* peptides, b) POH*^n^* complex with 3 Ang-(1-7)*^n^* peptides, c) PNH*^a^* in complex with 3 Ang-(1-7)*^n^* peptides and b) POH*^a^* at in complex with 3 Ang-(1-7)*^n^* peptides.

**Table 5:** Main interactions (HBs) formed between Ang-(1-7) peptide and PNH*^n^*. Only occupancies above 35% are shown. SegA correspond to dendrimer and SegB-SegD to peptides.

| Donnor | Acceptor | Occupancy (%) |
| --- | --- | --- |
| SegC-TYR4-Side-OH | SegA-INTR1-Main-O | 53.39 |
| SegA-TER1-Main-N | SegB-PRO7-Side-O2 | 52.70 |
| SegA-TER1-Main-N | SegD-PRO7-Side-O1 | 50.30 |
| SegA-TER1-Main-N | SegC-PRO7-Side-O1 | 49.73 |
| SegA-TER1-Main-N | SegC-PRO7-Side-O2 | 48.61 |
| SegA-TER1-Main-N | SegB-PRO7-Side-O1 | 48.60 |
| SegA-TER1-Main-N | SegD-PRO7-Side-O2 | 48.37 |
| SegA-TER1-Main-N | SegB-ASP1-Side-OD1 | 45.86 |
| SegA-TER1-Main-N | SegB-ASP1-Side-OD2 | 45.23 |
| SegC-ASP1-Main-N | SegA-CORE1-Main-O | 41.77 |
| SegA-INTR1-Main-N | SegC-TYR4-Side-OH | 39.49 |

### 4.3 Complexes stability

#### 4.3.1 PAMAM-NH/Ang-(1-7) complex

To identify global structural changes due to complex formation, the RMSD and the R*_g_* at bound state were measured and compared with that of the free structures. It can be seen in Figure 14ac that the binding of peptides stabilizes the dendrimer structure according to RMSD values. Meanwhile, as can be seen in Figure 14bd, the radius of gyration of the dendrimer and the peptides are not importantly affected by the binding.

**Figure 14:**
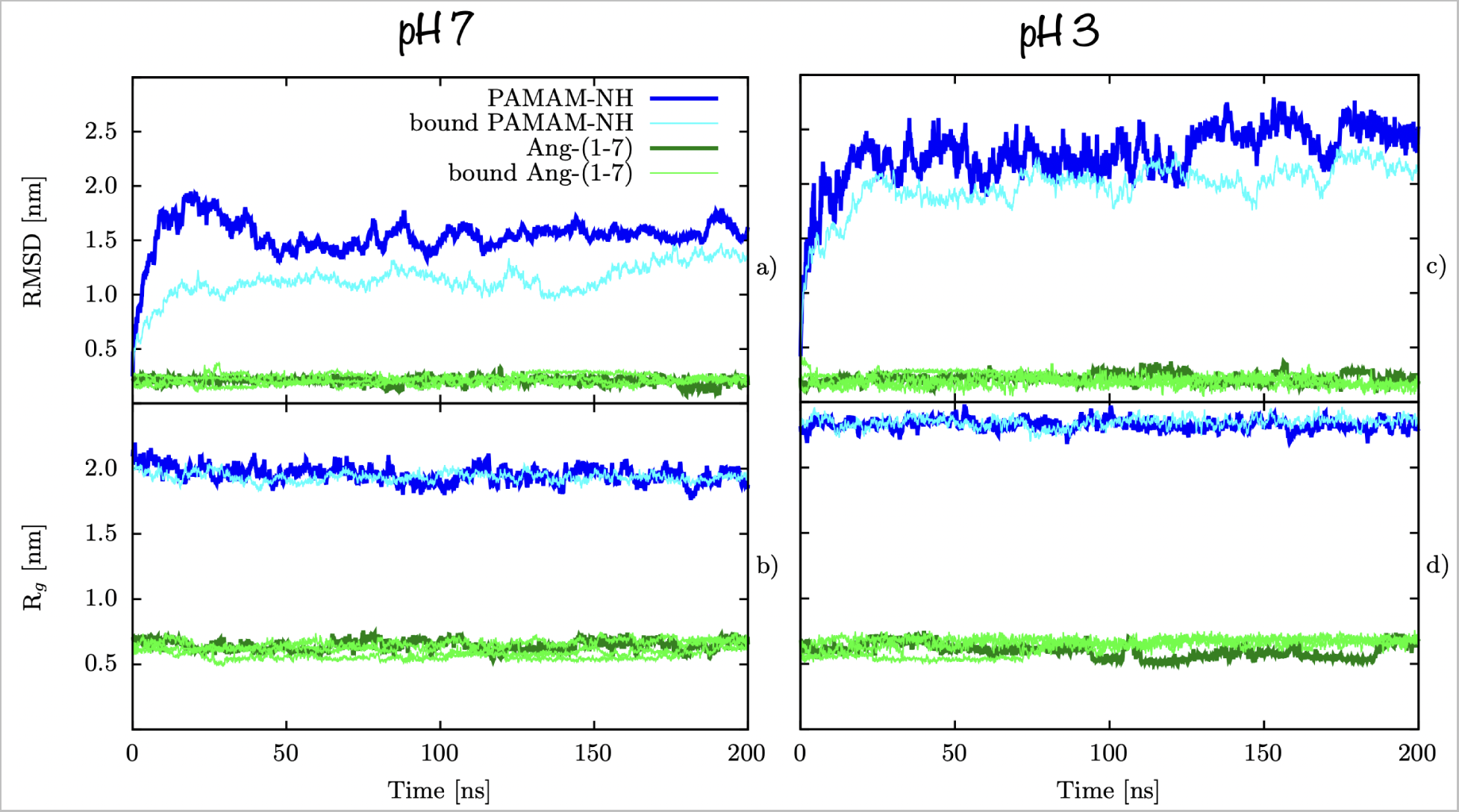
Structural stability changes of peptide or PAMAM-NH upon binding. RMSD of structures in complex compared with same structures free in solvent for a) neutral pH and c) acidic pH. R*_g_* of structures in complex compared with same structures free in solvent at b) neutral pH and d) acidic pH.

To test complex stability, the distance between the center of mass (COM) of the dendrimer and each peptide was measured. As showed in Figure 15a and Figure 18a, under neutral pH conditions, 2 peptides remained stable bonded during the simulation in the dendrimer internal domain around a distance of 1 nm and a third peptide remained stable bonded at the dendrimer surface around a distance of 3 nm, peptides remain mainly in the same initial sites. The dendrimer kept an open hourglass geometrical shape during the whole simulation time, allowing at least two peptides to be near the core. As shown in Figure 15d, under acidic pH conditions, from the 3 peptides initially at around 1 nm in the dendrimer internal domain, one remained very close to the core during all the simulation time, two went out quickly and stayed near the superficial zone, even leaving the dendrimer at certain points. From the two peptides in the dendrimer surface, one went to the back of the core at the last 50 ns of the simulation, interacting with the other peptide in the core. It appears that for two peptides, transitions from in to out the dendrimer had low energetic barriers or was an unstable state, in such a way that it is easy to get out, meanwhile, the third peptide appears to be in a stable deep minimum that forces it to stay close to the dendrimer core, but further research is required to confirm this hypothesis, due to the system flexibility, a MTD simulation could give us more insight about this process. Whith these results we can conclude that only one peptide remains stably bonded at the core of the dendrimer at acidic pH during the whole MD simulation.

**Figure 15:**
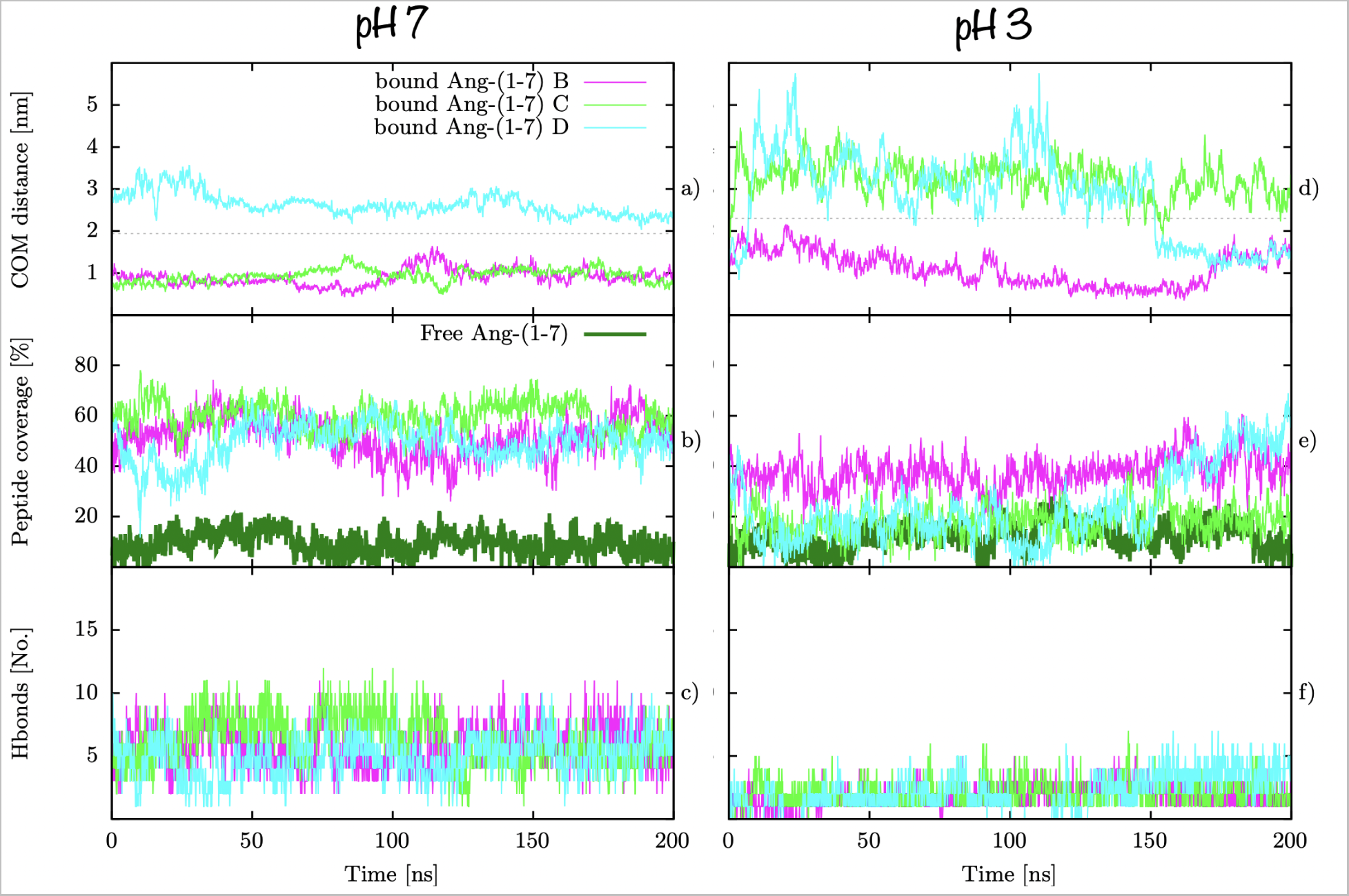
[PNH-A]*^n^*) stability. a) Distance from dendrimer COM to Ang-(1-7) peptides, b) peptide coverage according to SASA values and c) number of hydrogen bonds between dendrimer and peptides. # B-D refers to the different Ang-(1-7) peptides bonded to dendrimers. Grey dashed line represents the R*_g_* of the dendrimer as a reference for dendrimer periphery.

To test the protection capabilities of each dendrimer, the percent coverage of peptides due to binding, was measured. The free peptide in solution was taken as a reference, representing a molecule that is 100 % accessible to the solvent. As presented in Figure 15b, in the case of the [PNH-A]*^n^*, an average of 54% of coverage was found, confirming PNH*^n^* drug protection capabilities. As presented in Figure 15e, in the case of the [PNH-A]*^a^*, an average of 20% of coverage was found, a much lower percentage than in the case of neutral pH. This is reasonable considering that 2 peptides are in the superficial zone during most of the simulation time and even leave the dendrimer at certain points. If we consider only the peptide that remains at the core, it still has less coverage percentage compared with the ones at the core at neutral pH, probably because due to the open structure at acidic pH, cavities are also deferentially hydrated. It could mean that the peptide might be a little more accessible to water but due to its localization inside the dendrimer, it is not necessarily accessible to peptides.

The peptide protection is especially relevant for bonds susceptible to hydrolytic attack by endopeptidasas, as schematized in Figure 17. Once formed, Ang-(1-7) is rapidly hydrolyzed, especially by ACE^52^ and dipeptidyl peptidase 3 (DPP3).^11^ Here, we found a 46 % of [PNH-A]*^n^* coverage on the region attacked by DPP3 and a 73 % of [PNH-A]*^n^* coverage on the region attacked by ACE. This result is important because the very short half-life of Ang-(1–7) in the circulation is primarily accounted for peptide metabolism by ACE.^53^

**Figure 16:**
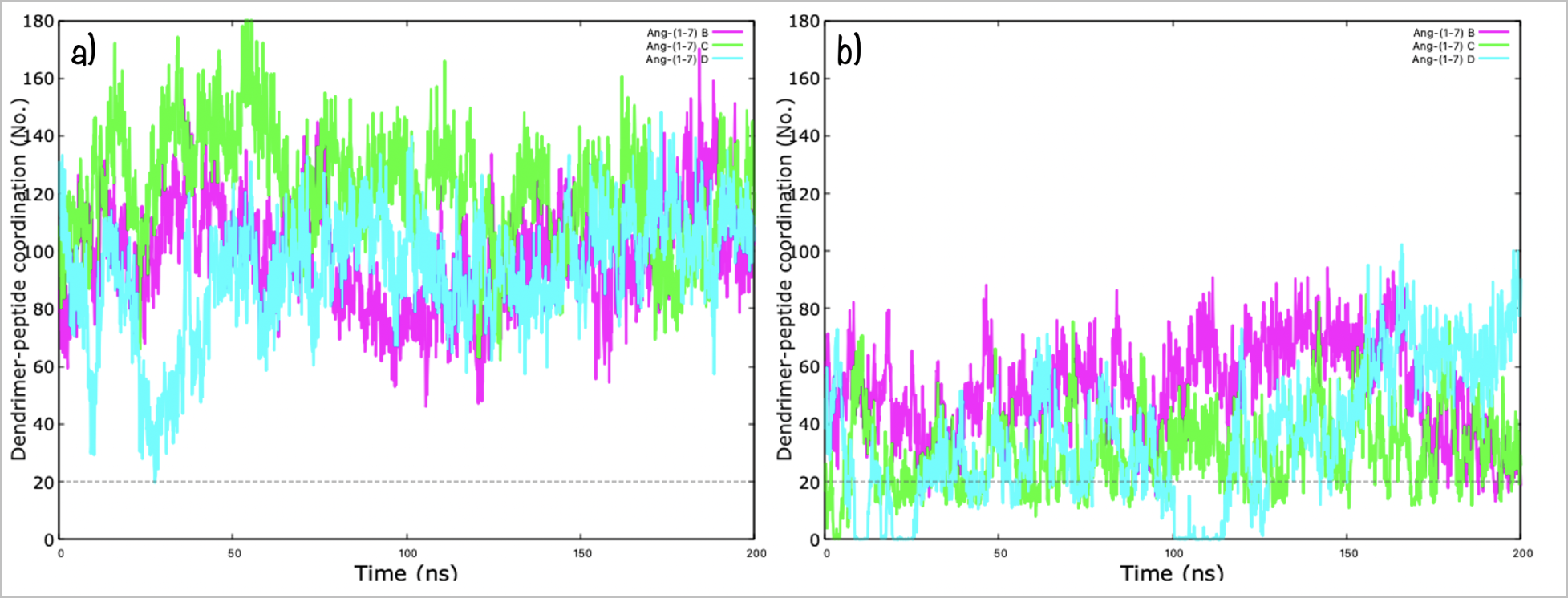
PAMAM-NH/Ang-(1-7) coordination number along simulation time at a) neutral pH and b) acidic pH. Coordination numbers are calculated that count the number of atoms from the second group that are within 0.3 nm of the first group. # B-D refers to the different Ang-(1-7) peptides bonded to dendrimers. Grey dashed line represents the threshold agreed that divides coupled from uncoupled states.

**Figure 17:**
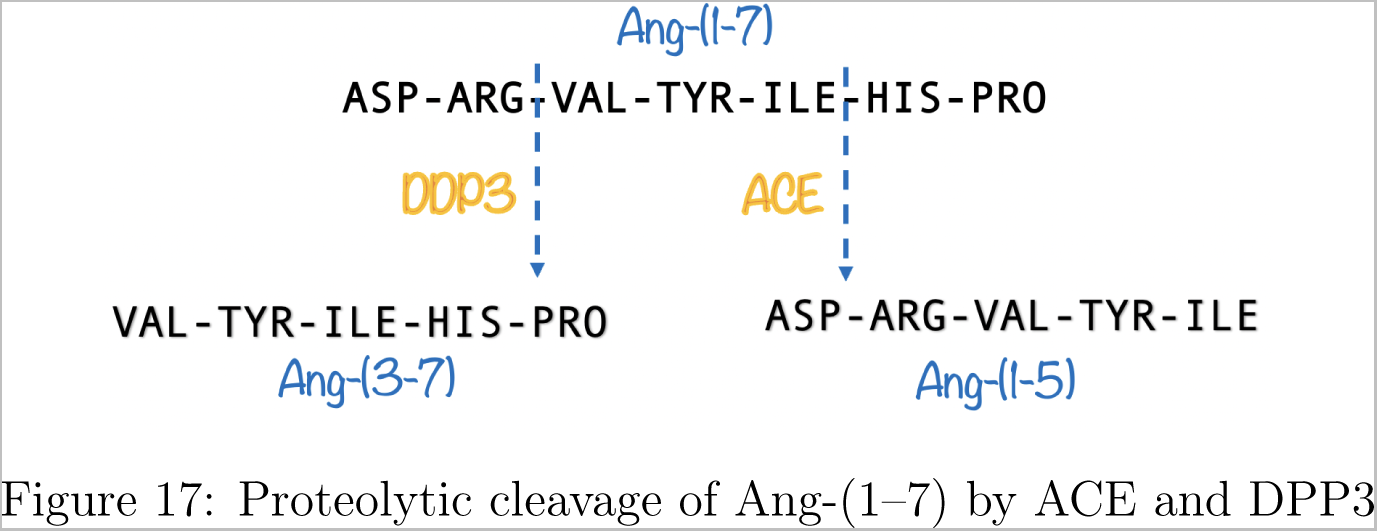
Proteolytic cleavage of Ang-(1–7) by ACE and DPP3

The main type of HBs formed between Ang-(1-7) and hydroxyl-terminated PAMAM dendrimer are presented in Figures S11 -S12, the number of NHBs through simulation time is presented in Figure 15cf and detailed occupancy in Tables 5 and 6. At neutral pH, a number of 5.4 HBs in average are formed between PNH*^n^* and Ang-(1-7) peptides, during the whole simulation time. The most populated HBs interactions are between core atoms, internal branches and terminal groups of PNH*^n^* with PRO7, ASP1 and TYR4 aminoacids in the peptide. Side chain of TYR4 in one of the peptides behaves as a donnor interacting with the amide Oxigen atom from the internal dendrimer as an acceptor during half of the simulation time; all the peptide keeps its interaction between its negative charged C-terminal and the main Nitrogen of the positively charged primary amine in the dendrimer terminal groups during almost half of the simulation time; ASP1 negative side chain interacts with the protonated primary amine of the dendrimer terminal groups during at least 40% of the time and also side chain of TYR4 interacts with the amide NH group from the internal dendrimer. The characteristic ASP1-TYR4/ASP1-VAL3 HB found in Ang-(1-7) metadynamics is still frequently formed (50 % of the simulation time) compared with its formation free in solvent (40 % of the simulation time), this implies that these intra peptide HBs are at least equally broken in favor of a peptide/dendrimer interaction (Table 5) compared with peptide/waters interaction. At acidic pH, a number of 2.5 HBs in average are formed between PNH*^a^* and Ang-(1-7) peptides, during the whole simulation time. The most populated HBs interactions are between internal branches of PNH*^n^* with ASP1 side chain in the peptide, but with much low occupations than in the case of neutral pH.

**Figure 18:**
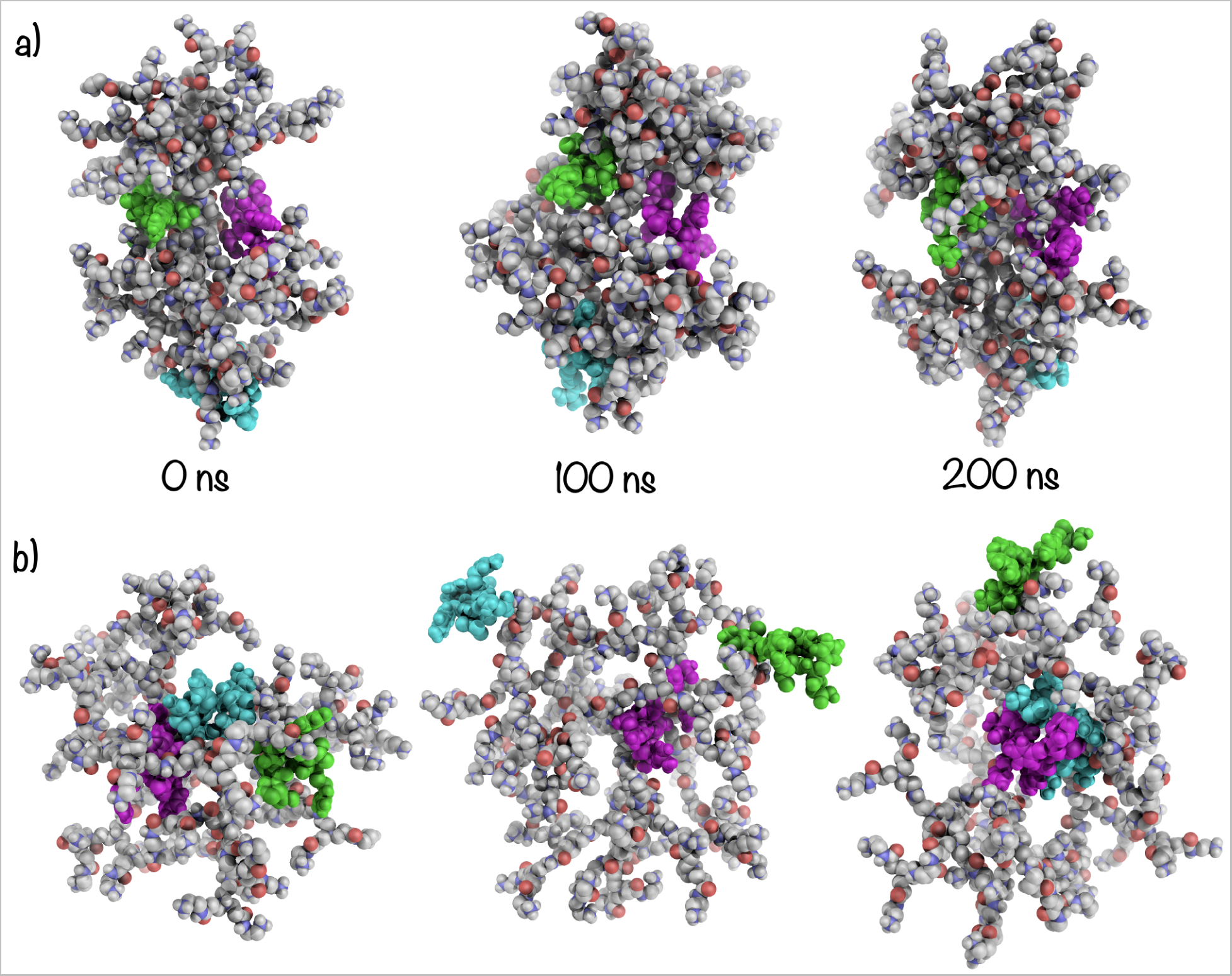
a) Time evolution of [PNH-A]*^n^* structure and b) occupancy percentage of the amino acids in the final 100 ns of MD simulation, determined by considering the contacts per amino acids at 3.0 Å from the dendrimer terminal groups (red bars) and to the dendrimer internal groups (blue bars). Occupancy denotes the average percentage of the three ang- (1–7) peptides that remained stable within the dendrimer.

**Table 6:**
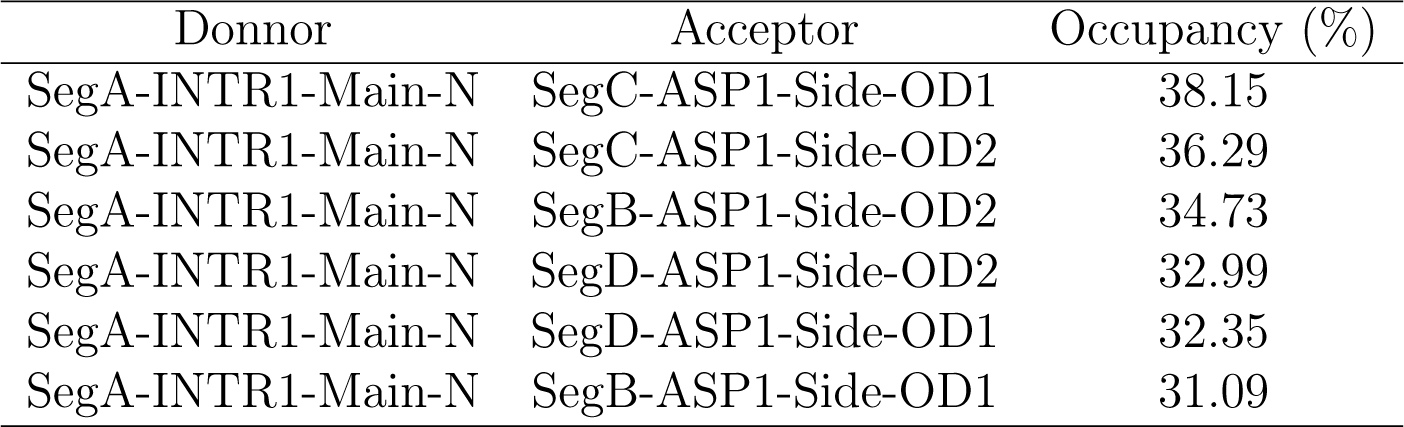
Main interactions (HBs) formed between Ang-(1-7) peptide and PNH*^a^*. Only occupancies above 10% are shown. SegA correspond to dendrimer and SegB-SegD to peptides.

| Donnor | Acceptor | Occupancy (%) |
| --- | --- | --- |
| SegA-INTR1-Main-N | SegC-ASP1-Side-OD1 | 38.15 |
| SegA-INTR1-Main-N | SegC-ASP1-Side-OD2 | 36.29 |
| SegA-INTR1-Main-N | SegB-ASP1-Side-OD2 | 34.73 |
| SegA-INTR1-Main-N | SegD-ASP1-Side-OD2 | 32.99 |
| SegA-INTR1-Main-N | SegD-ASP1-Side-OD1 | 32.35 |
| SegA-INTR1-Main-N | SegB-ASP1-Side-OD1 | 31.09 |

**Table 7:**
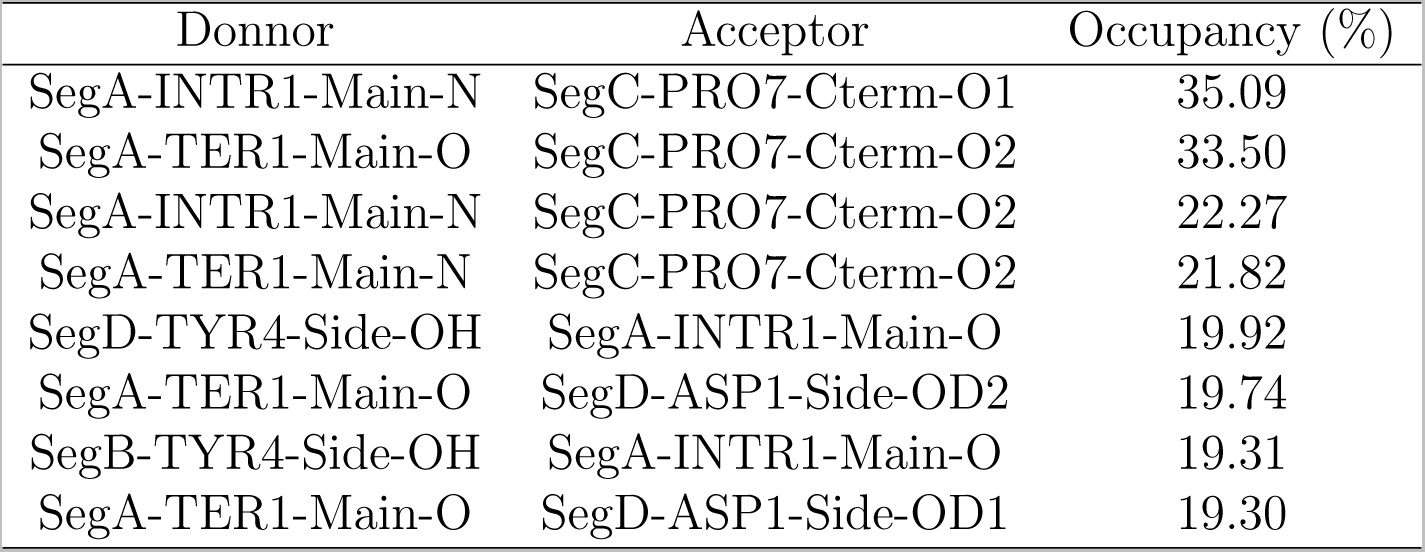
Main interactions (HBs) found between Ang-(1-7) peptide and POH*^n^* at neutral pH. Only occupancies above 19% are shown.SegA correspond to dendrimer and SegB-SegD with peptides.

According to Figure 19a, at neutral pH, TYR4 and ILE5 residues, interacted preferably with internal dendrimer groups rather than with the terminal groups, which was expected due to its hydrophobic/partially hydrophobic nature. On the other hand, ASP1 and PRO7 residues, interacted preferably with terminal dendrimer groups rather than with the internal groups, which was expected due to its negative charged side-chain/C-terminal groups. ARG2, VAL3 and HIS6 interacts similarly with terminal or internal dendrimer groups but in a less frequent way. As expected, arginine residue was more exposed to solvent due to its charged aminoacid side chain. According to Figure 19b, at acidic pH, ASP1, ARG2, TYR4 and HIS6 keep in contact preferably with internal dendrimer groups. Due to a more open dendrimer cavities, ASP1 and ARG2 are able to enter and interact with internal polar groups compared with the neutral case. Due to its hydrophobic/partially hydrophobic nature TYR4 keeps in contact preferably with internal dendrimer as in the neutral case but less frequently. HIS6 at acidic pH is considered as positively charged, while at neutral pH is neutral, this change modify its preference to be in contact with internal dendrimer groups instead with positively charged terminal groups. Finally, VAL3, ILE5 and PRO7 hydrophobic residues keep almost no contact with the dendrimer, here C-terminal is neutral compared with the neutral case. Taking into account the number of peptides stably bonded near the core, the higher percent coverage of peptides from water, the major number of HBs between dendrimer and peptides during the whole simulation time, the frequency in which this HBs occur and the peptide and dendrimer frequency of contacts, it is likely that amino terminated PAMAM dendrimer can act as a better encapsulator near neutral pH and as a release agent under acidic pH. However, further studies are needed in this direction.

**Figure 19:**
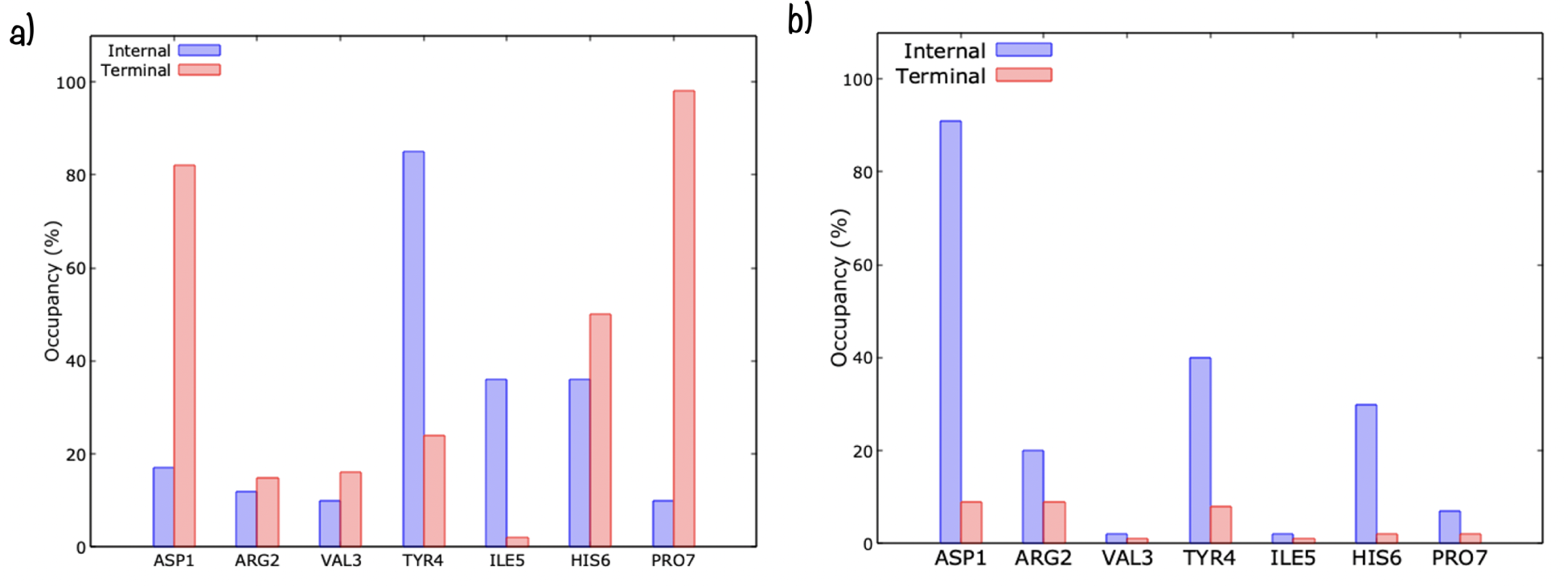
a) Time evolution of [PNH-A]*^n^* structure and b) occupancy percentage of the amino acids in the final 100 ns of MD simulation, determined by considering the contacts per amino acids at 3.0 Å from the dendrimer terminal groups (red bars) and to the dendrimer internal groups (blue bars). Occupancy denotes the average percentage of the three ang- (1–7) peptides that remained stable within the dendrimer.

#### 4.3.2 PAMAM-OH/Ang-(1-7) complex

We measured RMSD as a kind of first indicator of stability upon complex formation. It can be seen in Figure 20ac that the binding of peptides slightly stabilizes the dendrimer structure at neutral pH but, it does not happen at acidic pH. Besides that, the structure of the peptides are not greatly affected by the binding to the dendrimer. For their part, the R*_g_* of either the dendrimer or the peptide are not importantly affected by the binding. Higher fluctuations on dendrimer RMSD and R*_g_* at acidic pH correspond to the continuous movement of the protonated groups which makes it difficult to be stabilized by ligand binding.

**Figure 20:**
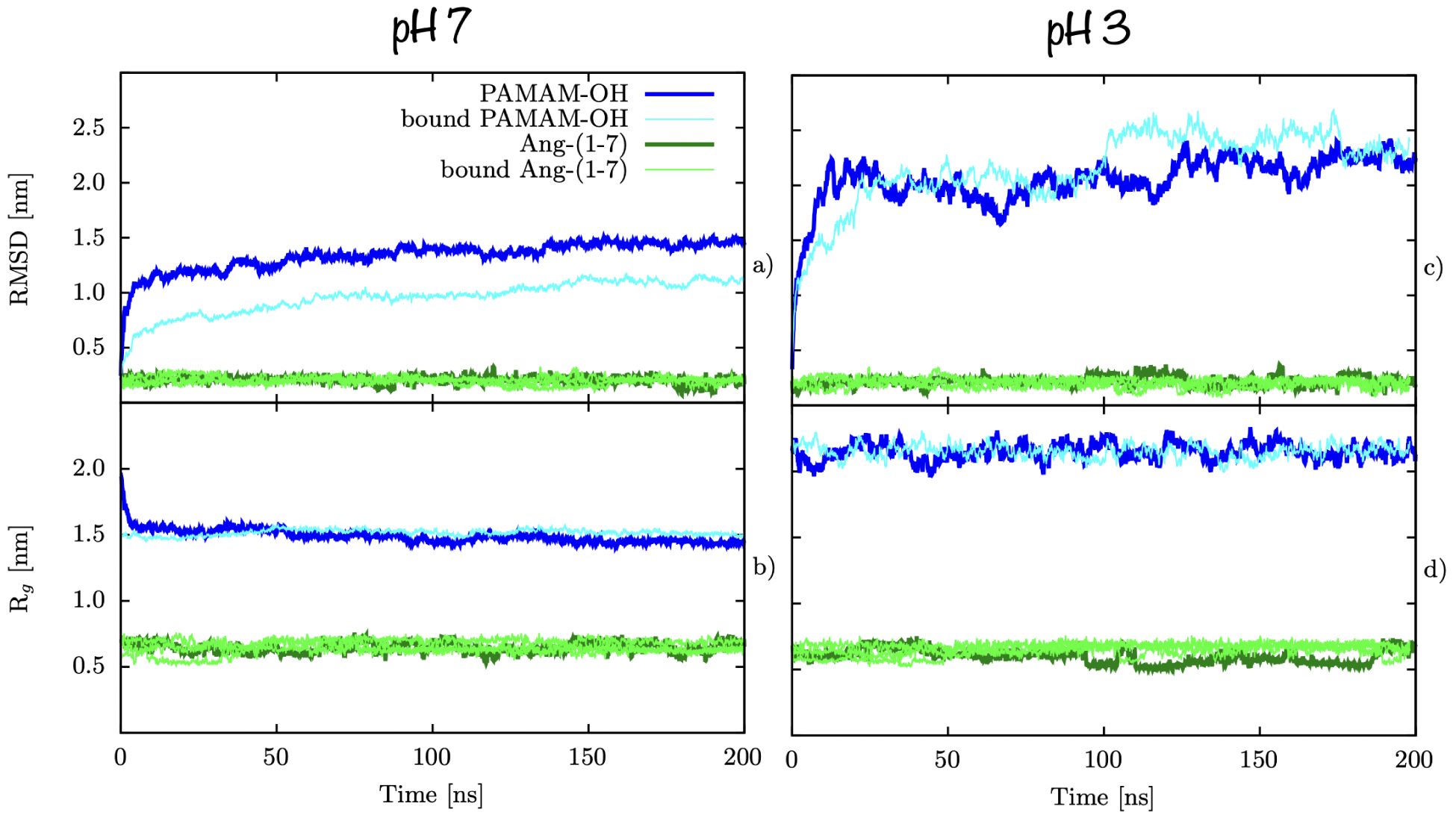
Structural stability changes of peptide or PAMAM-OH upon binding. RMSD of structures in complex compared with same structures free in solvent for a) neutral pH and c) acidic pH. R*_g_* of structures in complex compared with same structures free in solvent at b) neutral pH and d) acidic pH.

To further evaluate complex stability, the distance between the Center Of Mass (COM) of the dendrimer and each peptide was measured. As shown in Figure 21a, under neutral pH conditions, all the three Ang-(1–7) peptides remained at a stable distance of 1.6 *±* 0.1 nm from the dendrimer COM, this is, mostly on the dendrimer surface taking into account that the R*_g_* is of 1.46 nm, as also illustrated in the timeline evolution of frames in Figure 23a. A closer view into the simulations shows that peptides in [POH-A]*^n^*, explore the nearby area around its initial binding site, which could be due to the symmetry of the dendrimer groups. Moreover, the spherical compacted shape of the dendrimer appears to avoid peptide’s complete internalization. The above results might imply that the energetic barriers between superficial binding sites are low, allowing the peptide to go from one local minimum to another, in contrast, the energy barrier needed to leave the dendrimer appears to be high enough, keeping the peptides interacting with the dendrimer as shown in Figure 22a, but further investigation is needed to confirm energetic landscape of the process. In the case of [POH-A]*^a^* in Figure 21d and Figure 23b, all the three peptides started at the internal dendrimer region around 1.2 nm, quickly both of them move into the periphery, around the R*_g_* of 2.3 nm, only transiently re-entering to the dendrimer interior. Conversely, the third peptide stays in the dendrimer interior and only transiently moves to the periphery. The number of contacs between the dendrimer and the peptides are presented in Figure 22b, were we can observe how the three peptides unbind and bind several times during the MD simulation.

**Figure 21:**
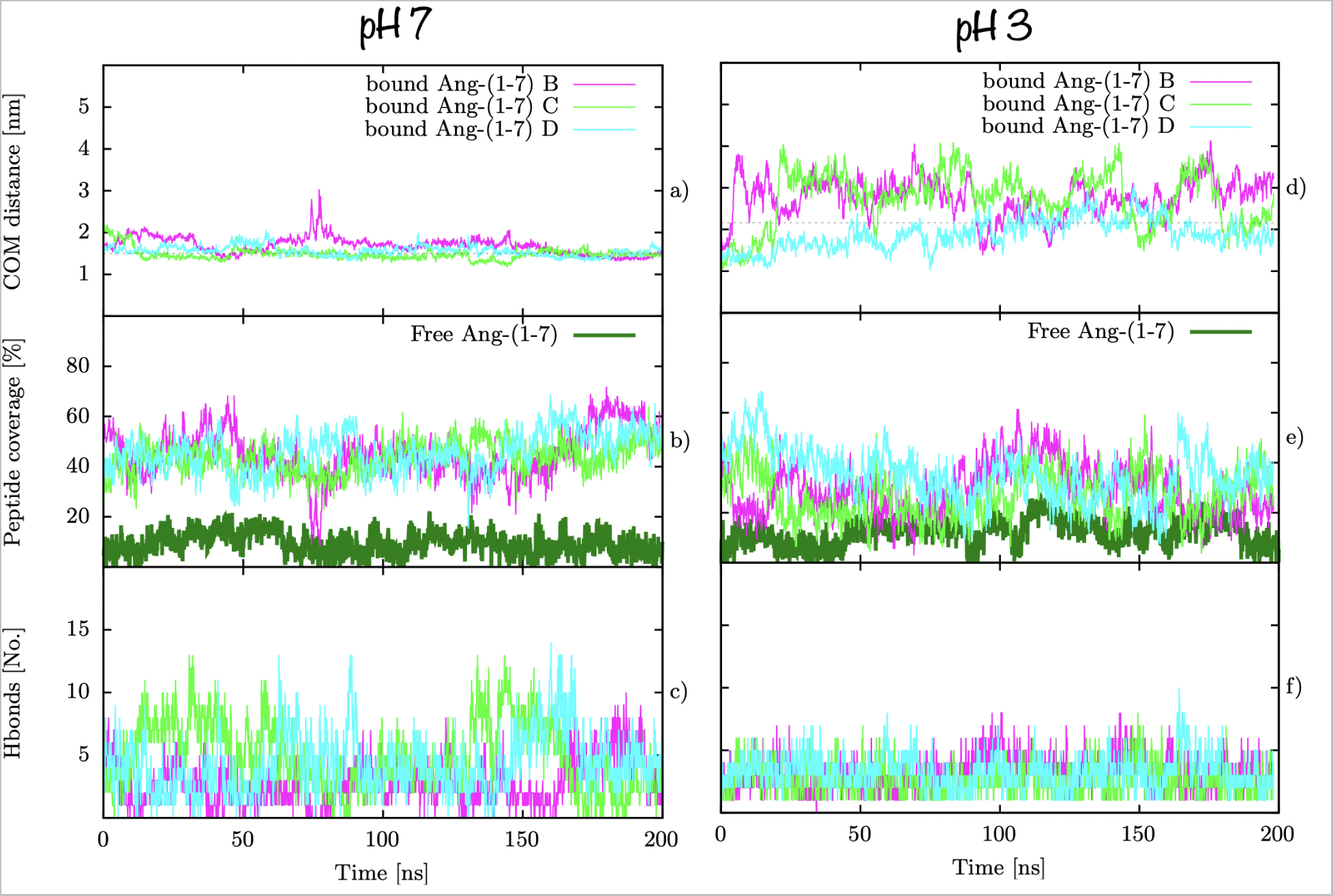
[POH-A]*^n^* stability. a) Distance from dendrimer COM to Ang-(1-7) peptides, b) peptide coverage according to SASA values and c) number of hydrogen bonds between dendrimer and peptides. # B-D refers to the different Ang-(1-7) peptides bonded to dendrimers. Grey dashed line represents the R*_g_* of the dendrimer as a reference for dendrimer periphery.

**Figure 22:**
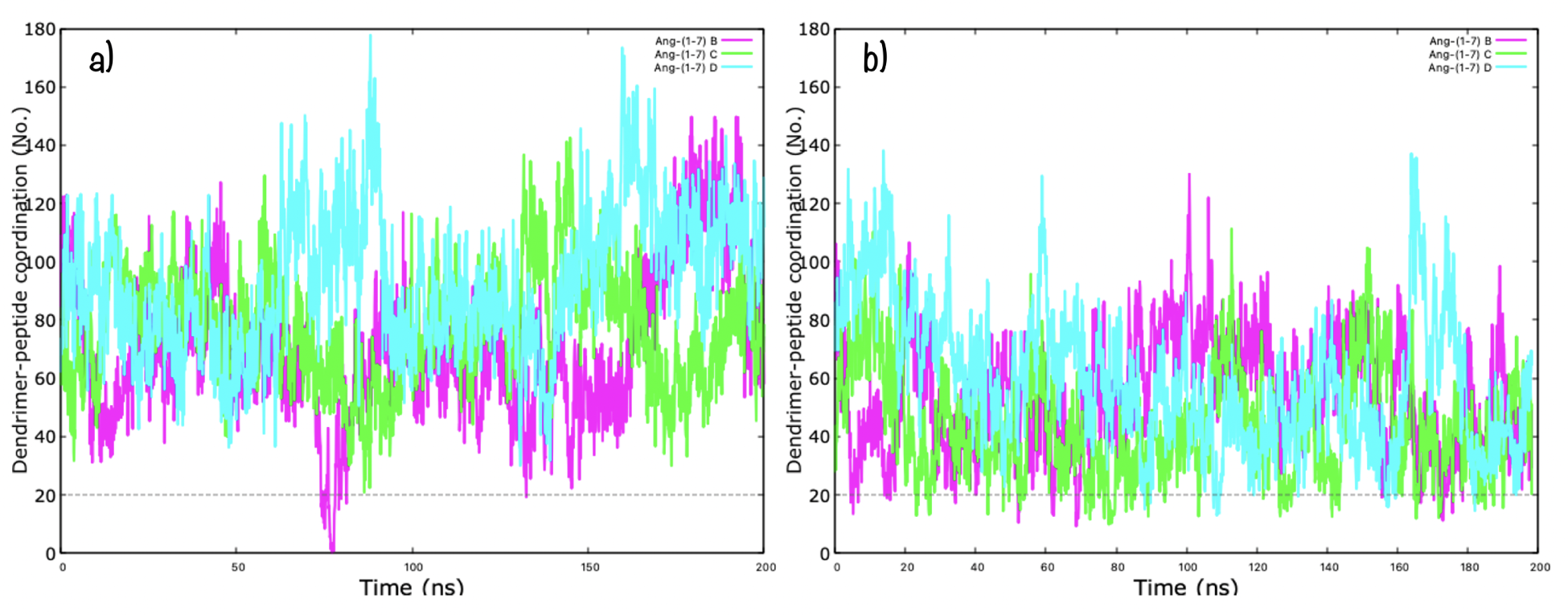
PAMAM-OH/Ang-(1-7) coordination number along simulation time at a) neutral pH and b) acidic pH. Coordination numbers are calculated that count the number of atoms from the second group that are within 0.3 nm of the first group. # B-D refers to the different Ang-(1-7) peptides bonded to dendrimers. Grey dashed line represents the threshold agreed that divides coupled from uncoupled states.

**Figure 23:**
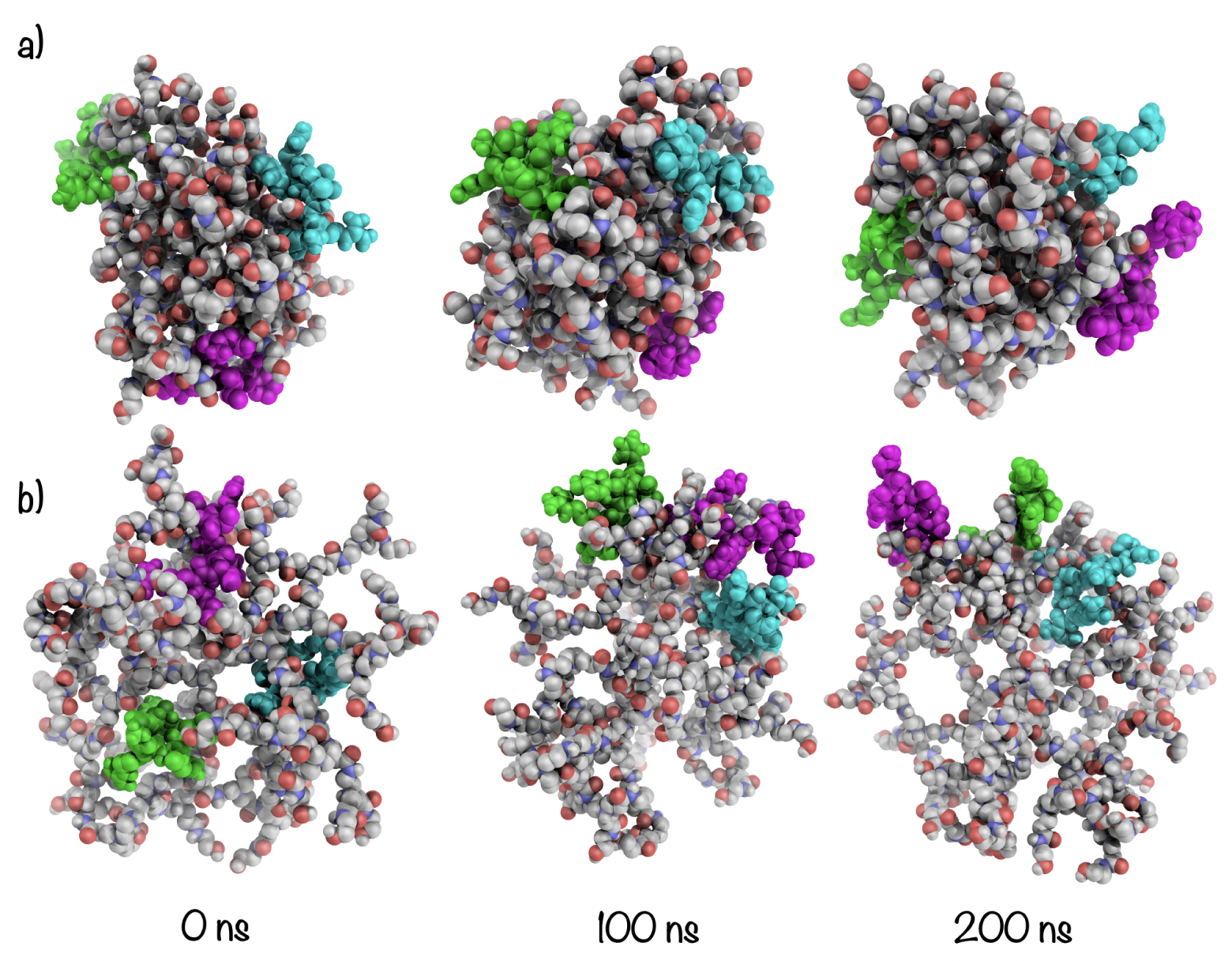
Time evolution of PAMAM-OH/Ang-(1-7) complex structures at a) neutral pH and b) acidic pH.

For testing the protection capabilities of each dendrimer, the percent coverage of peptides due to binding, was measured. As presented in Figure 21b, in the case of [POH-A]*^n^* protection, the peptides showed a dendrimer coverage of around 44%. As presented in Figure 21e, in the case of [POH-A]*^a^* protection, the peptides showed a dendrimer coverage of around 30%, lesser coverage than in the case of neutral pH. At neutral pH, where peptides coverage is bigguer, we found a 55 % of [POH-A]*^n^* coverture on the region attacked by DPP3 and a 57 % of [POH-A]*^n^* coverture on the region attacked by ACE.

The main type of HBs formed between Ang-(1-7) and amino-terminated PAMAM dendrimer are presented in Figures S13 -S14, the number of hydrogen bonds (NHBs) through simulation time is presented in Figure 21cf and detailed occupancy in Tables 7 and 8. At neutral pH, a number of 4 HBs in average are formed between POH*^n^* and Ang-(1-7) peptides, during the whole simulation time. It was found until 144 different HBs between POH*^n^* and peptides, however they have low % of occupancy (below 36 %), suggesting a low specificity of these bonds. The most populated HBs interactions are between internal branches and terminal groups of POH*^n^* with PRO7, ASP1 and TYR4 aminoacids in the peptide, no interactions with core are present above 19 % of occupancy. At acidic pH, a number of 3 HBs in average are formed between POH*^a^* and Ang-(1-7) peptides, during the whole MD simulation time. It was found until 139 different HBs between POH*^n^* and peptides, however, only ASP1 and TYR4 with internal branches have considerable % of occupancy. The most populated HBs interactions are between internal branches of POH*^n^* with charged side chain of ASP1 and backbone of TYR4 in the peptide, no interactions with core and terminal groups are present above 19 % of occupancy. At neutral pH, the characteristic ASP1-TYR4/ASP1-VAL3 HB found in Ang-(1-7) metadynamics is still frequently formed (53 % of the simulation time) compared with its formation free in solvent (40 % of the simulation time), this imply that these intra peptide HBs are at least equally broken but this is not necessarily in favor of a peptide/dendrimer interaction (Table 7), it is also because part of the time the residue is solved exposed and interacts with waters.

**Table 8:** Main interactions (HBs) formed between Ang-(1-7) peptide and POH*^a^*. Only occupancies above 19% are shown. SegA correspond to dendrimer and SegB-SegD to peptides.

| Donnor | Acceptor | Occupancy (%) |
| --- | --- | --- |
| SegA-INTR1-Main-N | SegD-ASP1-Side-OD1 | 50.88 |
| SegA-INTR1-Main-N | SegD-ASP1-Side-OD2 | 49.80 |
| SegA-INTR1-Main-N | SegB-ASP1-Side-OD1 | 40.68 |
| SegA-INTR1-Main-N | SegB-ASP1-Side-OD2 | 40.39 |
| SegA-INTR1-Main-N | SegC-ASP1-Side-OD1 | 39.36 |
| SegA-INTR1-Main-N | SegC-ASP1-Side-OD2 | 39.32 |
| SegB-TYR4-Main-N | SegA-INTR1-Main-O | 19.88 |

According to Figure 24a, at neutral pH, TYR4 interacts preferably with internal dendrimer groups rather than with the terminal groups, which was expected due to its partially hydrophobic aromatic nature which prefers to be buried in a hydrophobic core and its polar hydroxyl (-OH) group also being able to interact with polar groups in the dendrimer. Particularly, the inclusion of the TYR residue is favored in both dendrimer types, as in the case of *β*-cyclodextrin complexes. ^14^ In the other hand, ASP1, ARG2, VAL3 and PRO7 residues, stay in contact slightly more with terminal dendrimer groups rather than with the internal groups, in case of negative charged ASP1 and C-terminal PRO7 with polar hydroxil terminal group in the dendrimer, arginine residue was in contact with terminal groups in the dendrimer and exposed to solvent due to its charged aminoacid side chain. ILE5 and HIS6 keep in contact similarly with terminal or internal dendrimer groups. At acidic pH, ASP1, ARG2, VAl3 and TYR4 interact preferibly with internal dendrimer groups. ILE5, HIS6 and PRO7 interact equally with internal and terminal groups but in a less frequent manner.

**Figure 24:**
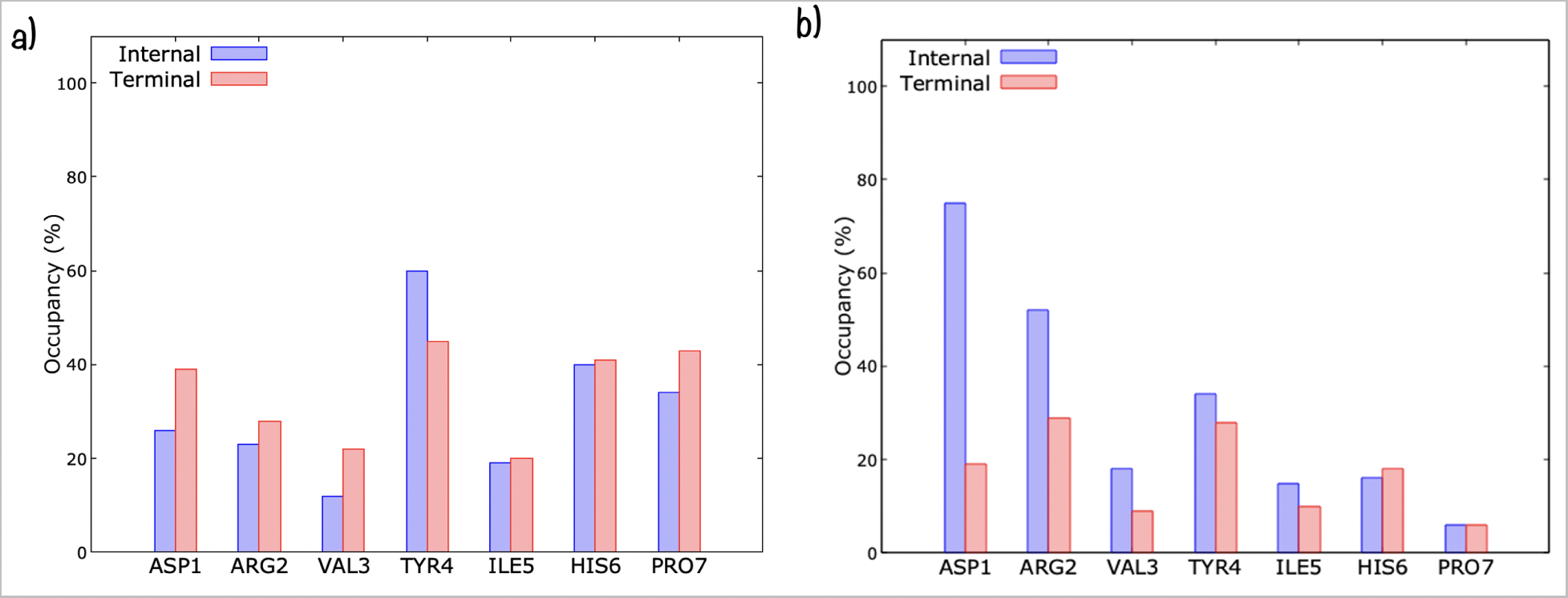
Occupancy percentage of the amino acids in the final 100 ns of MD simulation at a) neutral pH and b) acidic pH. Determined by considering the contacts per amino acids at 3.0 Å from the dendrimer terminal groups (red bars) and to the dendrimer internal groups (blue bars). Occupancy denotes the average percentage of the three ang-(1–7) peptides that remained stable within the dendrimer.

Taking into account the number of peptides stable bonded at the surface, the higher percent coverage of peptides from water, the slightly major number of HBs between dendrimer and peptides during the whole simulation time, the frequency in which this HBs occur and the peptide and dendrimer frequency of contacts, it is likely that hydroxyl terminated PAMAM dendrimer can act as a Ang-(1-7) carrier at neutral pH and as a release agent under acidic pH. However, further experimental studies are needed in this direction.

So far, it appears that POH*^n^* is able to interact with 3 peptides but not necessarily encapsulate them, however, this is valid for the case when dendrimer and peptide are prepared at neutral pH separately and afterwads put togheter. It remains still the question if the encapsulation is reached under different conditions, what happen with the complex when pH becomes neutral? In order to address this question, we designed a new complex (complex 2) by using an open dendrimer structure (from the non-equilibrated region of the dendrimer in solution), two peptides were placed in the internal cavities by our double-docking approach. This complex was evolved in an MD during 200 ns, evolution of the peptides is showed in Figure S7. As can be seen, initially, peptides are close from the core in the internal voids of the dendrimer, eventually the dendrimer becomes more and more compact and finally, the peptides remain interacting at the dendrimer surface. This give us an idea of what could happen if the encapsulation is previously done, the complex at neutral pH evolve to a situation similar to the presented previusly (Figure 24a), dendrimer prefers a compact structure and peptides interact mostly with external surface. This is the case of reference,^21^ where dendrimer initial open structure for complex formation appears to be taken from a non-equilibrated structure, eventually dendrimer becomes more and more compact, even releasing one of the peptides, but at the end of the 60 ns MD productions, simulations does not necessarily reach convergence and could show a meta-stable state where peptides are partially encapsulated. Here, we show that in the last 100 ns of the MD production, most part of the peptides are interating with external dendrimer parts whether it started with the peptides in or out, with slightly differences. Details are showed in Figure S8.

## 5 Conclusions

Structural parameters such as the radius of gyration, the asphericity and the radial distribution function confirmed that the GROMOS-compatible 2016H66 force field, together with our chosen methodology are capable to model the theoretical and experimental structural features of PAMAM dendrimers evaluated in this work.

The classical MD simulations of PAMAM dendrimers in solution showed that hydroxyl-terminated PAMAM dendrimer at neutral pH has a more compact, less structured, more spherical, less hydrated and less flexible structure than amino-terminated PAMAM dendrimer. These results emphasises how a single change in terminal group could lead to differential behaviour of dendrimer structure and eventually in Ang-(1-7) binding. Also, this structural information is envisioned to prove useful for the encapsulation of other drugs.

The classical MD simulations of Ang-(1-7) in solution yielded to conformations in which a bend secondary structure is stabilized by interactions between residues VAL3 and TYR4, which is in agreement with NMR experimental studies. It was also found that at acidic pH, 60 % of the Ang-(1-7) conformations were similar to the NMR solution structure, meanwhile, at neutral pH, this also happened but in a slightly less frequent percentage of conformations.

In the accelerated MD metadynamics simulations, Ang-(1-7) free energy surface showed a unique broad free energy minimum on the radius of gyration and donor-acceptor contact phase space, confirming the Ang-(1-7) flexibility due to its peptide nature. One dimensional free energy surface as a function of radius of gyration showed that free energy barriers between metastable states in the main basin are on the order of 1 kT, thus easily passed.

At neutral pH, PAMAM dendrimers with both terminal types are able to interact with at least 3 peptides in a stable way, with a coverage of up to 70%. However, hydroxyl-terminated PAMAM dendrimer only partially encapsulate the peptides due to its compact structure.

At acidic pH, PAMAM dendrimers with both terminal groups are still able to interact with positevely charged Ang-(1-7) peptides, either internalized or in its periphery, however, contacts, coverage and HBs are lesser than at neutral pH, suggesting a state for peptide release.

With the results found so far, it appears that amino-terminated PAMAM dendrimer at neutral pH posses slightly better features to load and protect Ang-(1-7) peptides than hydroxyl-terminated PAMAM dendrimer. However, either hydroxyl-terminated PAMAM dendrimer as well as amino-terminated PAMAM dendrimer are able to bind Ang-(1-7) peptides in a differential form, suggesting that hydroxyl-terminated PAMAM dendrimer is able to act as nanocarrier/nanotransporter more than nanoprotector of this peptide.

For the PAMAMNH(OH)/Ang-(1-7) complexes, the therapeutic mechanism could be encapsulate or capture the peptide at near neutral pH and deliver them at acidic pH, where repulsions between protonated primary and/or tertiary amines give rise to a open and hydrated environment, allowing positively charged peptides to be solvated and released easily.

Finally, we propose that the complexes studied here should be experimentally tested to validate the properties predicted through *In Silico* approaches, as potential therapeutic agents.

