## Supplemental Material File for "Unveiling the G4-PAMAM capacity to bind and protect Ang-(1-7) bioactive peptide"

#### 1 Peptide: structure and protonation states.

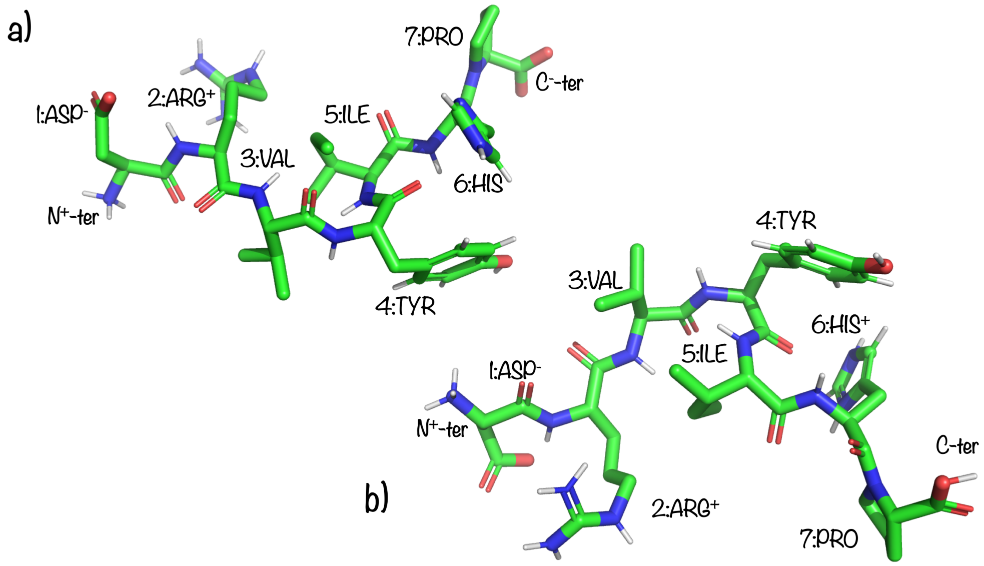


Figure S1: Illustration of the initial structure used for Ang-(1-7) at a) neutral pH and b) acidic pH.

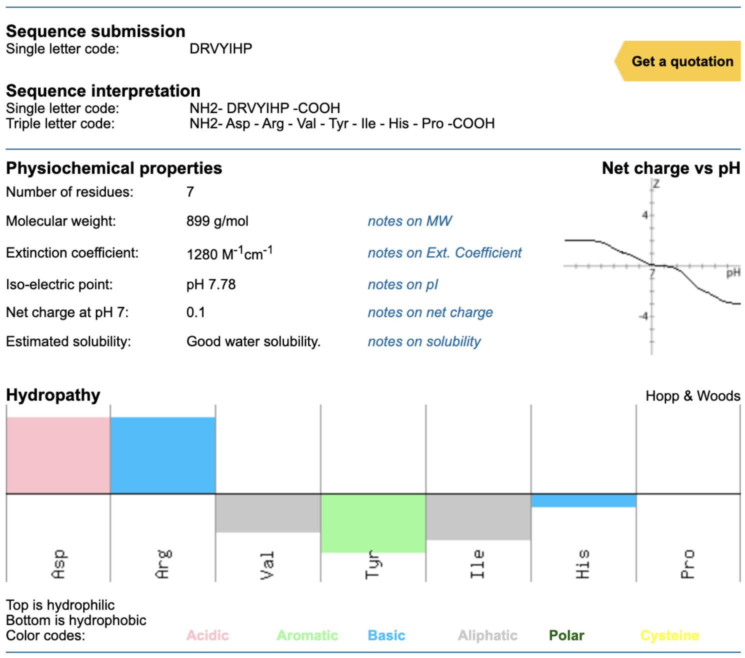


Figure S2: Peptide physicochemical properties according to INNOVAGEN peptide property calculator.

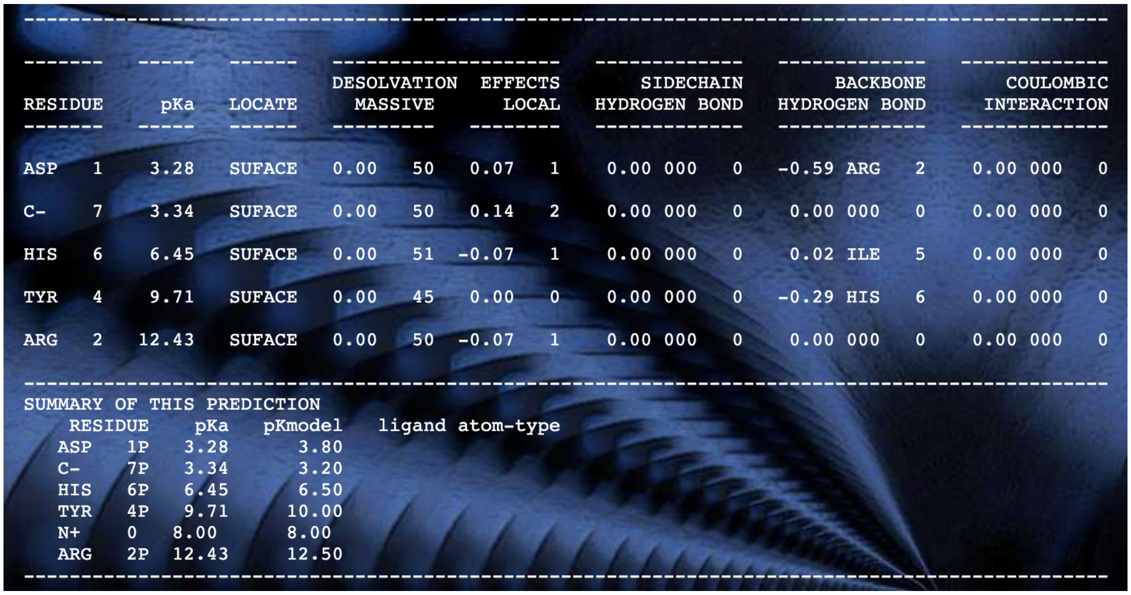

| RESIDUE | | pKa | LOCATE | DESOLVATION MASSIVE | | EFFECTS LOCAL | | SIDECHAIN HYDROGEN BOND | | | BACKBONE HYDROGEN BOND | | | COULOMBIC INTERACTION | | |
|---|---|---|---|---|---|---|---|---|---|---|---|---|---|---|---|---|
| ASP | 1 | 3.28 | SUFACE | 0.00 | 50 | 0.07 | 1 | 0.00 | 000 | 0 | -0.59 | ARG | 2 | 0.00 | 000 | 0 |
| C- | 7 | 3.34 | SUFACE | 0.00 | 50 | 0.14 | 2 | 0.00 | 000 | 0 | 0.00 | 000 | 0 | 0.00 | 000 | 0 |
| HIS | 6 | 6.45 | SUFACE | 0.00 | 51 | -0.07 | 1 | 0.00 | 000 | 0 | 0.02 | ILE | 5 | 0.00 | 000 | 0 |
| TYR | 4 | 9.71 | SUFACE | 0.00 | 45 | 0.00 | 0 | 0.00 | 000 | 0 | -0.29 | HIS | 6 | 0.00 | 000 | 0 |
| ARG | 2 | 12.43 | SUFACE | 0.00 | 50 | -0.07 | 1 | 0.00 | 000 | 0 | 0.00 | 000 | 0 | 0.00 | 000 | 0 |

SUMMARY OF THIS PREDICTION

| RESIDUE | | pKa | pKmodel | ligand atom-type |
|---|---|---|---|---|
| ASP | 1P | 3.28 | 3.80 | |
| C- | 7P | 3.34 | 3.20 | |
| HIS | 6P | 6.45 | 6.50 | |
| TYR | 4P | 9.71 | 10.00 | |
| N+ | 0 | 8.00 | 8.00 | |
| ARG | 2P | 12.43 | 12.50 | |

Figure S3: Peptide pKa values according to PropKa program.

#### 2 Dendrimers: structure, protonation states and parameters.

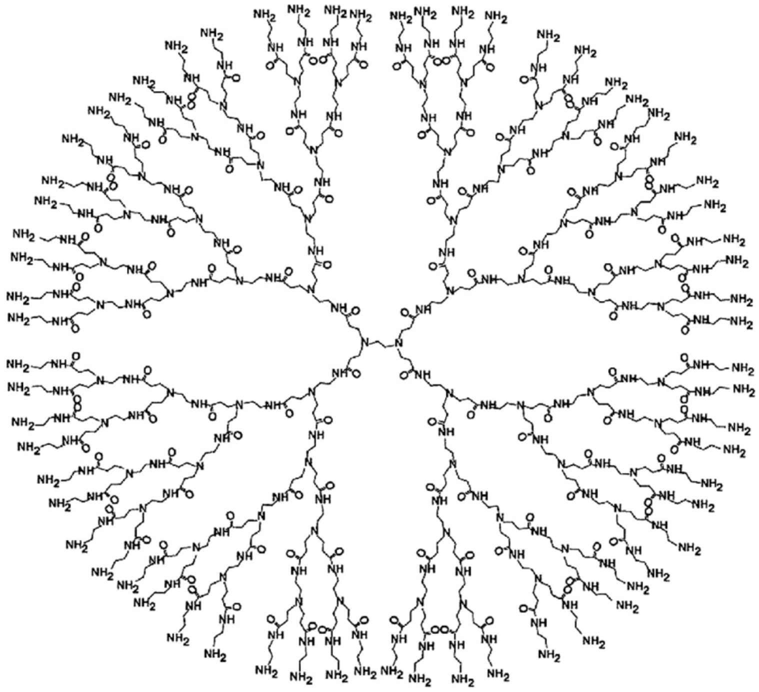

Figure S4: PAMAM-$NH_2$ G4 chemical structure.

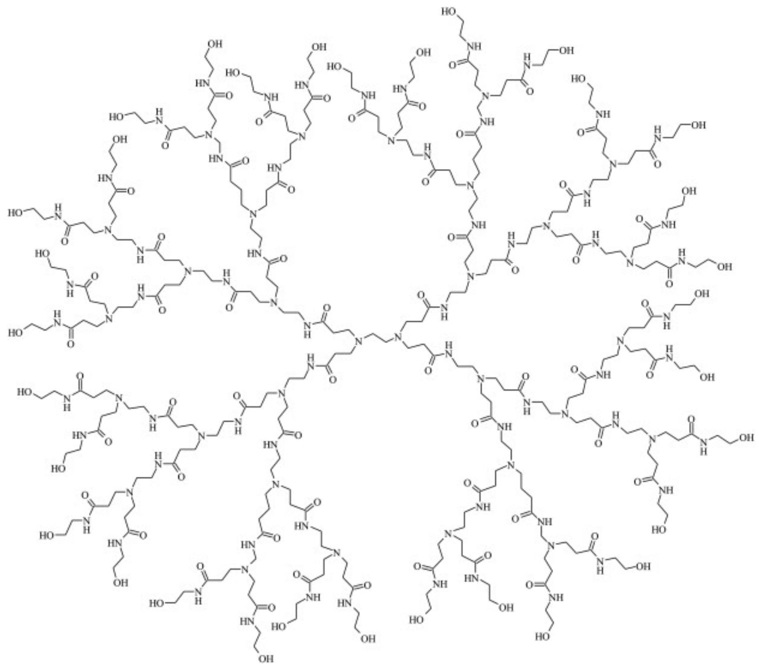

Figure S5: PAMAM-OH G4 chemical structure.

Table S1: Building-block definition for the PAMAM-OH terminal group used in the present work. The columns correspond to the chemical structure, atom numbering (ID), atom type, partial atomic charge (q) and atomic mass. The asterisk in the chemical structure indicates an open valency for connecting to either the core group or a monomer group.

| Chemical structure | Atom ID | Atom Type | q | mass |
|---|---|---|---|---|
| 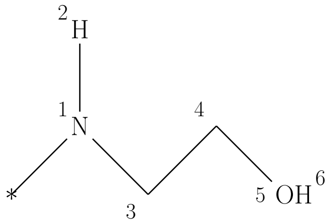 | 1 | N | -0.500 | 14.00670 |
| | 2 | H | 0.310 | 1.00800 |
| | 3 | CH2 | 0.190 | 14.02700 |
| | 4 | CH2 | 0.290 | 14.02700 |
| | 5 | OA | -0.700 | 15.99940 |
| | 6 | H | 0.410 | 1.00800 |

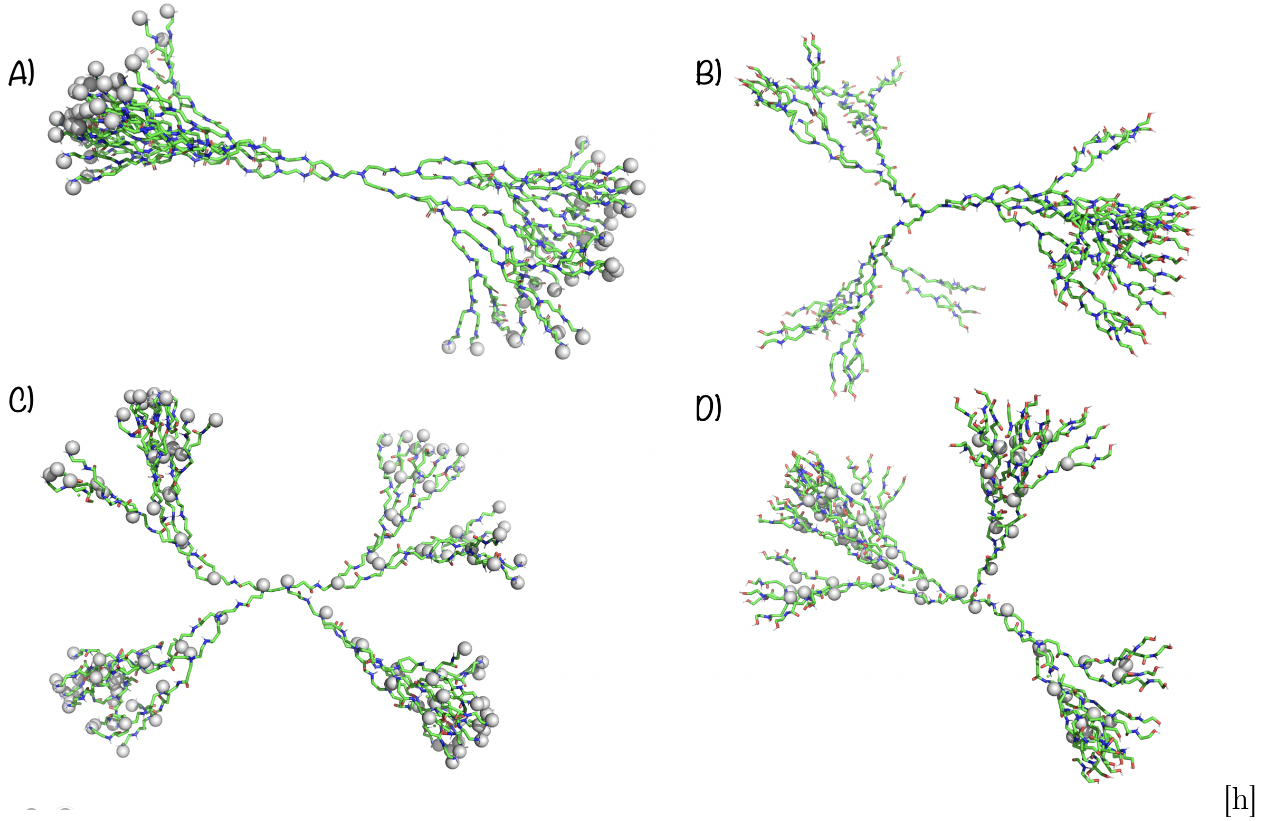


[h]

Figure S6: Illustration of the protonation states considered in the present work for the PAMAM dendrimer: A) PAMAM-$NH_2$ at neutral pH has its primary amines protonated and B) PAMAM-OH at neutral pH has no protonations. Structures correspond to the ones generated by Pypolybuilder. The additional hydrogen atoms at the protonated sites are displayed as the gray VDW spheres.

### 3 Double docking approach

Table S2: Characteristics of docking results. Dendrimer conformer (d) and peptide conformer (p).

| conformers | PAMAM-$NH_2^n$ Affinity [kcal/mol] | HBs | PAMAM-$OH^n$ Affinity [kcal/mol] | HBs | PAMAM-$NH_2^a$ Affinity [kcal/mol] | HBs | PAMAM-$OH^a$ Affinity [kcal/mol] | HBs |
|---|---|---|---|---|---|---|---|---|
| d0p0 | -4.8 | 8 | -5.8 | 7 | -4.2 | 12 | -6.7 | 11 |
| d0p1 | -5.4 | 6 | -4.7 | 7 | -5.8 | 7 | -6.0 | 19 |
| d0p2 | -5.5 | 5 | -5.2 | 6 | -4.6 | 9 | -5.7 | 16 |
| d0p3 | -5.3 | 6 | -5.6 | 7 | 4.9 | 13 | -5.8 | 21 |
| d0p4 | -5.3 | 7 | -4.5 | 5 | 6.0 | 14 | -6.5 | 16 |
| d0p5 | -6.3 | 5 | -4.8 | 2 | -4.8 | 13 | -6.0 | 12 |
| d1p0 | -6.3 | 10 | -5.2 | 7 | -4.7 | 20 | -5.1 | 20 |
| d1p1 | -5.9 | 6 | -5.4 | 8 | -4.3 | 4 | -5.3 | 16 |
| d1p2 | -6.3 | 11 | -5.0 | 8 | -4.3 | 20 | -5.6 | 19 |
| d1p3 | -6.4 | 10 | -6.0 | 9 | -4.6 | 17 | -5.1 | 8 |
| d1p4 | -6.4 | 8 | -5.4 | 6 | -5.1 | 17 | -5.7 | 20 |
| d1p5 | -6.3 | 8 | -5.0 | 3 | -4.4 | 19 | -4.9 | 7 |
| d2p0 | -5.6 | 6 | -6.8 | 4 | -6.1 | 17 | -6.2 | 16 |
| d2p1 | -4.6 | 4 | -5.5 | 8 | -3.7 | 3 | -4.0 | 4 |
| d2p2 | -5.1 | 8 | -6.2 | 5 | -5.3 | 26 | -4.7 | 6 |
| d2p3 | -6.2 | 7 | -5.8 | 4 | -5.6 | 28 | -3.9 | 8 |
| d2p4 | -5.3 | 8 | -6.0 | 3 | -3.8 | 5 | -6.0 | 12 |
| d2p5 | -5.9 | 9 | -6.1 | 10 | -5.2 | 21 | -6.3 | 22 |

### 4 PAMAM-OH/Ang-(1-7) complex 2

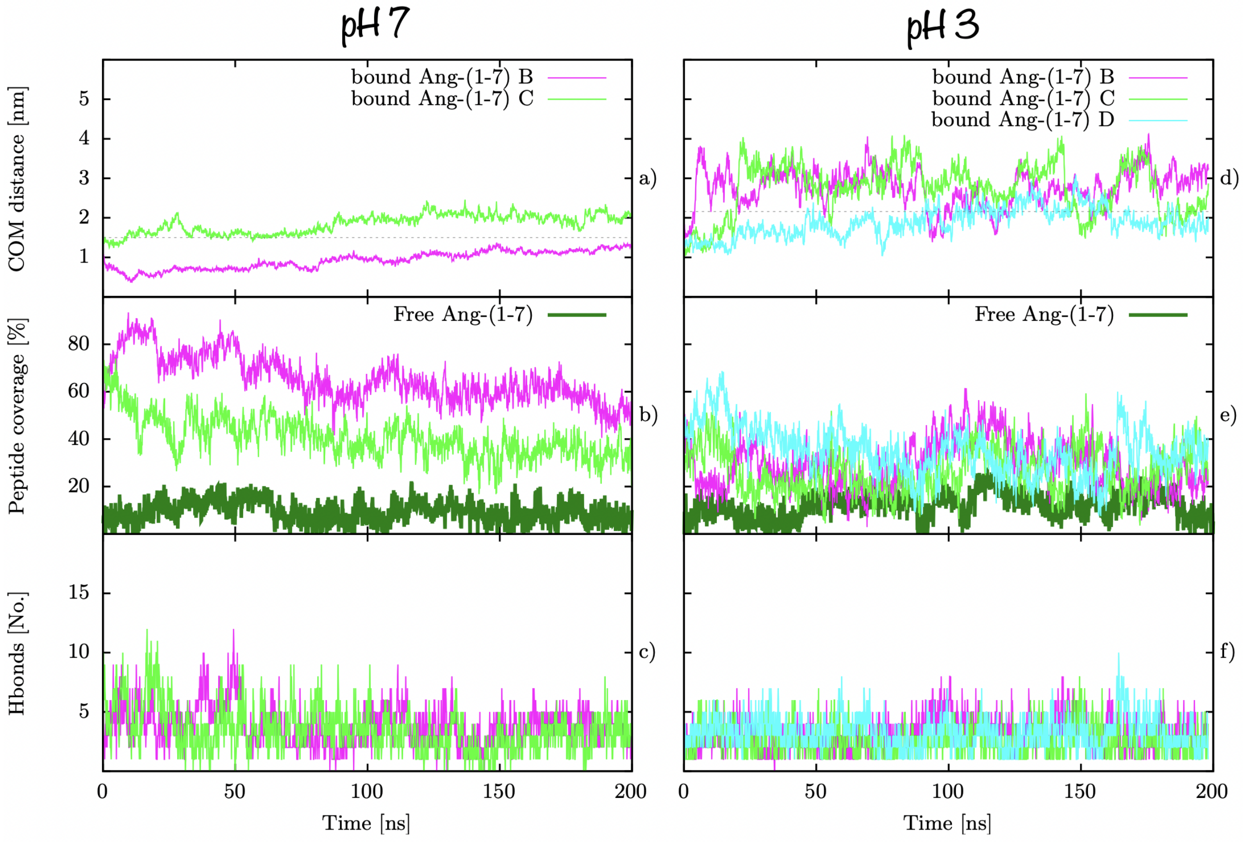


Figure S7: PAMAM-OH/Ang-(1-7) complex 2 stability. a) Distance from dendrimer COM to Ang-(1-7) peptides, b) peptide coverage according to SASA values and c) number of hydrogen bonds between dendrimer and peptides. # B-D refers to the different Ang-(1-7) peptides bonded to dendrimers. Grey dashed line represents the $R_g$ of the dendrimer as a reference for dendrimer periphery.

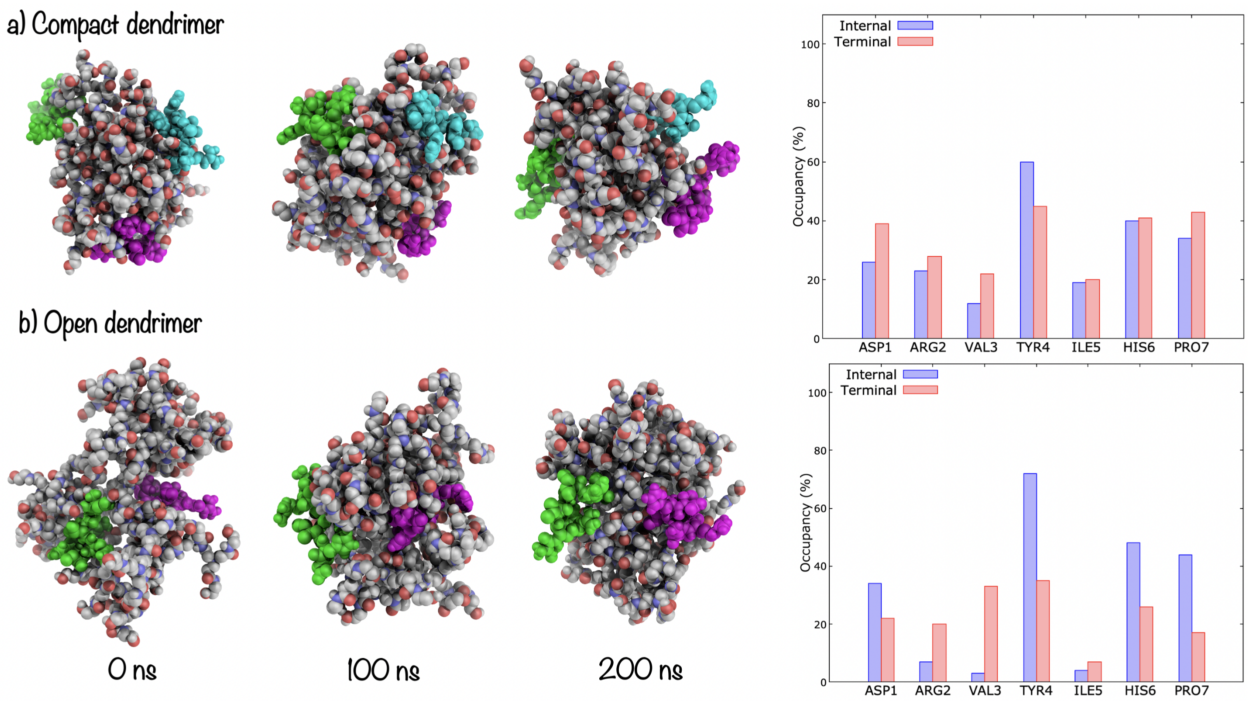


Figure S8: $[POH\text{-}A]^n$ complex 2 time line evolution of structures and occupancy.

### 5 Metadynamics

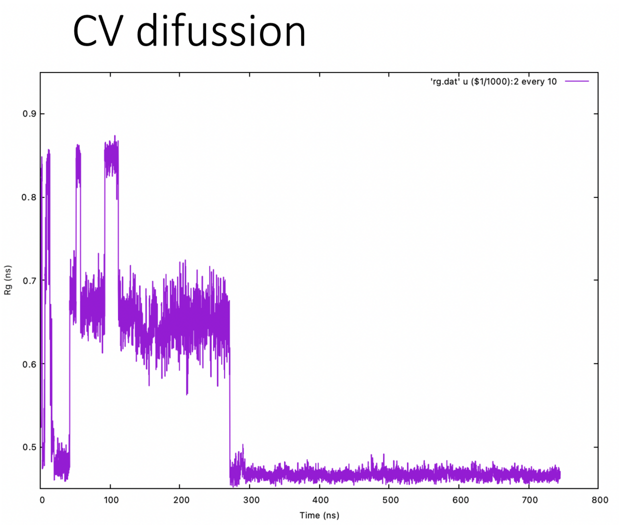


Figure S9: Metadynamics of Ang-(1-7) when implementing only 1CV ($R_g$). $R_g$ diffusion as a function of simulation time.

### 6 Ang-(1-7) hydrogen bonds detail

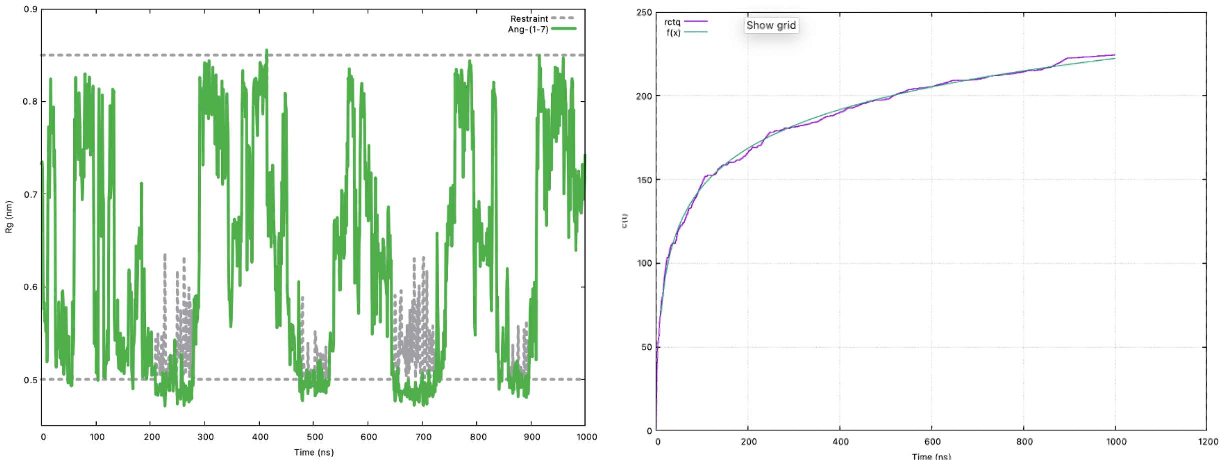


Figure S10: Metadynamics of Ang-(1-7) in function of 2 CVs ($R_g$ and $C_{d-a}$). a) Time evolution of the $R_g$) CV and b) C(t) fitted to a logaritmic model as a function of simulation time. Restraint is showed as grey dashed line.

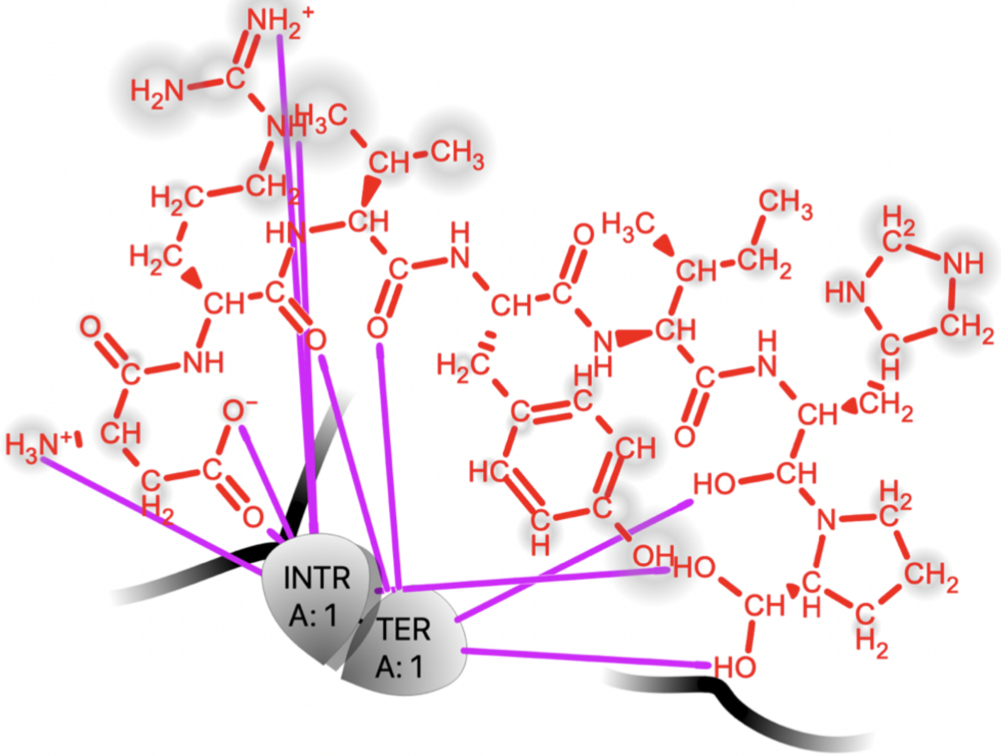


| Acceptor | Donor | |
|---|---|---|
| ASP1-Side-(C-O) | PAMAMOH-N-TER-(N-H) | (Atom numbers 1278- N 164) |
| ASP1-Side-(C=O) | PAMAMOH-TER-(N-H) | (Atom numbers 1277- N 999) |
| PAMAMOH-N-TER-(C-O) | ARG2(N-H) | (Atom numbers N1287-O784) |
| PAMAMOH-TER-(C-O) | ASP1(N-H3) | (Atom number N1270-O1135) |
| PAMAMOH-N-TER-(C-O) | ARG2(N-H) | (Atom numbers N1293-O784) |
| ARG2-side(C=O) | PAMAMOH-TER-(N-H) | (Atom numbers N993-O1297) |
| PRO7-side(C-O) | PAMAMOH-N-TER-(N-H) | (Atom number O1353-N99) |
| PRO7-side(C-O) | PAMAMOH-TER-(N-H) | (Atom numbers O1354-N927) |
| HISB6-side(C-O) | PAMAMOH-TER-(N-H) | (Atom numbers O1345-N1171) |
| VAL3-side(C=O) | PAMAMOH-TER-(N-H) | (Atom numbers O1305-N993) |

Figure S11: Details on the type of hydrogen bonds that are able to be formed between Ang-(1-7) and PAMAMOH in complex at neutral pH.

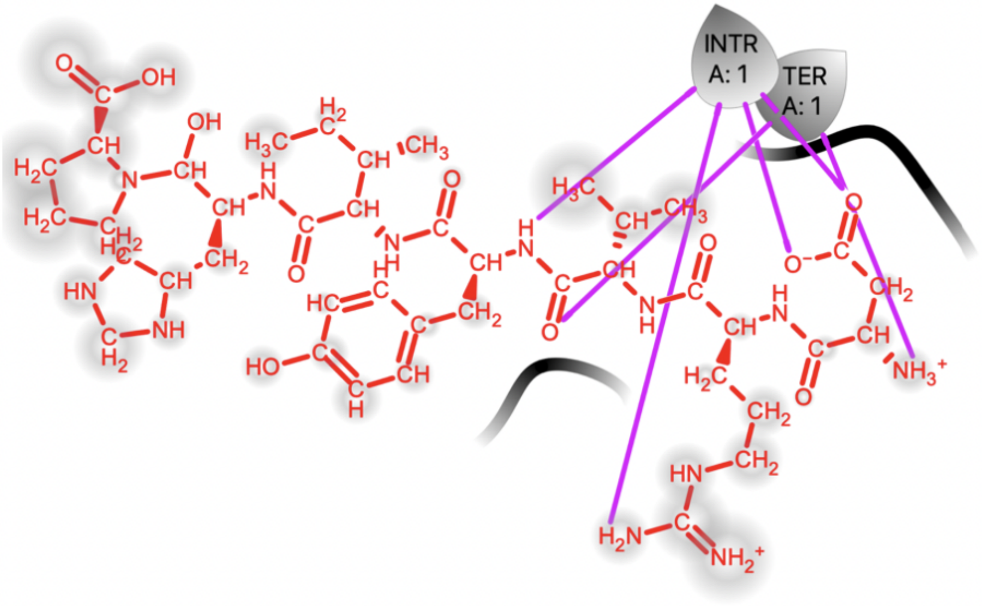


| Acceptor | Donor | |
|---|---|---|
| ASP1-Side-(C=O) | PAMAMOH-N-TER-(N-H) | (Atom numbers 1428- N 107) |
| ASP1-Side-(C-O) | PAMAMOH-N-TER-(N-H) | (Atom numbers 1429- N 111) |
| PAMAMOH-TER-(C-O) | ASP1(NH3) | (Atom numbers N1421-O999) |
| PAMAMOH-N-TER-(C-O) | ARG2(N-H) | (Atom number N1441-O624) |
| PAMAMOH-TER-(OH) | VAL3(C=O) | (Atom numbers N1456-O1042) |
| PAMAMOH-N-TER-(C-O) | TYR4(N-H) | (Atom numbers N1457-O117) |

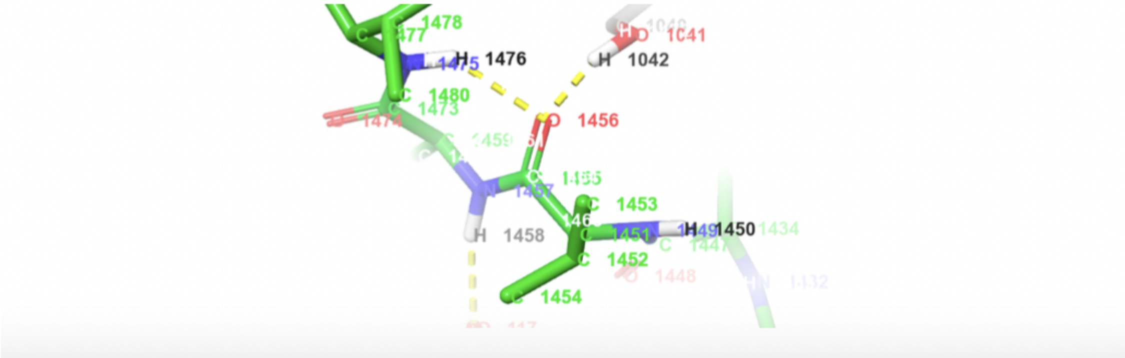


Figure S12: Details on the type of hydrogen bonds that are able to be formed between Ang-(1-7) and PAMAMOH in complex at acidic pH.

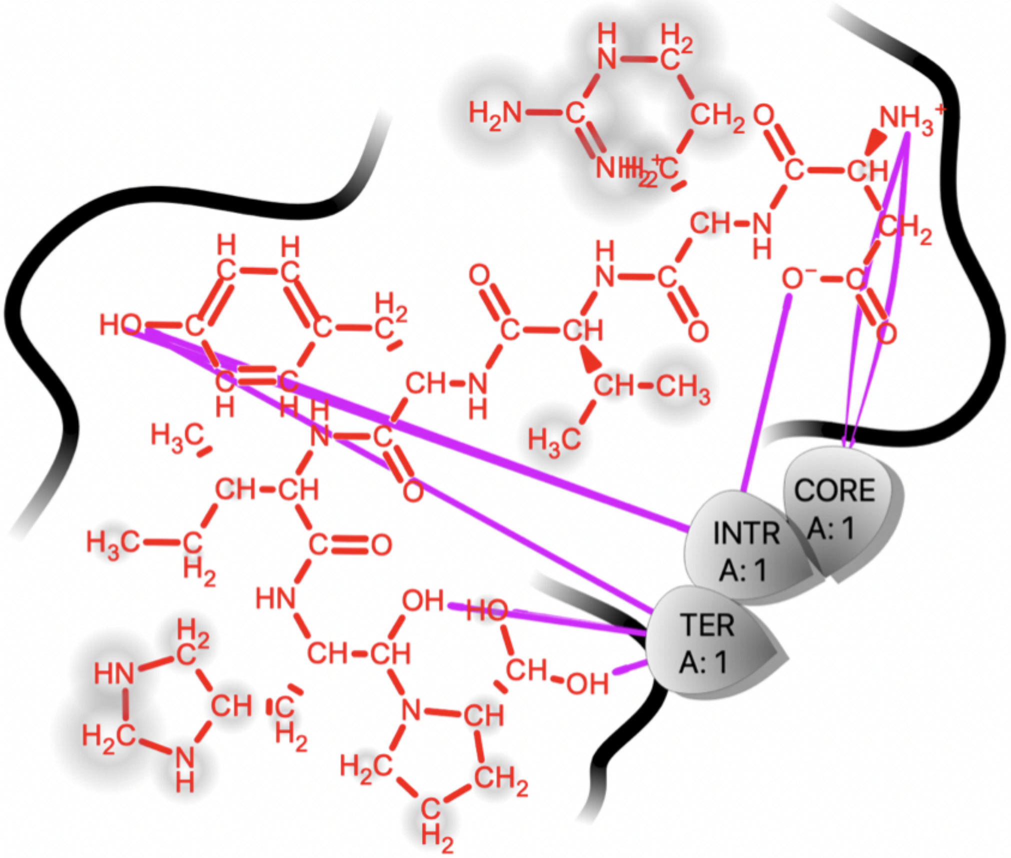


| Acceptor | Donor | |
|---|---|---|
| PRO7-Side-(C-O) | PAMAMNH-TER-(N-H) | (Atom numbers O1481- N 1233) |
| PRO7-Side-(C-O) | PAMAMNH-TER-(NH3) | (Atom numbers O1482- N 845) |
| HISB6-Side-(C-O) | PAMAMNH-TER-(NH3) | (Atom numbers O1474- N 845) |
| TYR4-side(O-H) | PAMAMNH-TER-(N-H) | (Atom numbers O1448-N 961) |
| ASP1(C-$O^-$) | PAMAMNH-TER-(NH3) | (Atom numbers O1406-N965) |
| PAMAMNH-N-TER-(C-O) | ASP1($NH3^+$) | (Atom number O443-N1398) |

Figure S13: Details on the type of hydrogen bonds that are able to be formed between Ang-(1-7) and PAMAMNH in complex at neutral pH.

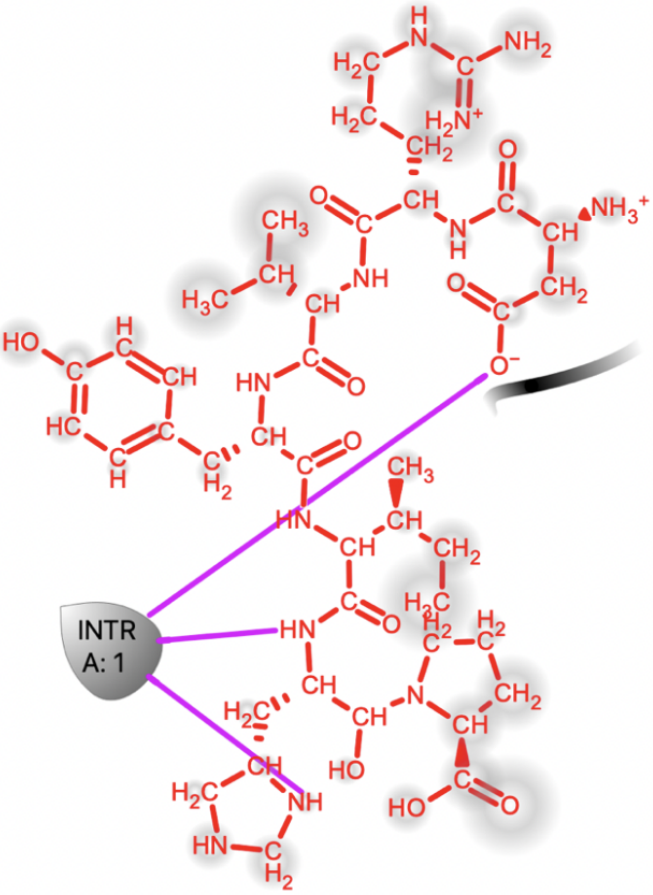


| Acceptor | Donor |
| --- | --- |
| PAMAMNH-N-TER-(C-O) | HISH6-Side-(NH) (Atom numbers N1617- O214) |
| PAMAMNH-N-TER-(C-O) | HISH6-Side-(NH) (Atom numbers N1612- O33) |
| ASP1-Side-(C-O$^-$) | PAMAMNH-N-TER-(N) (Atom numbers O1557- N671 845) |

Figure S14: Details on the type of hydrogen bonds that are able to be formed between Ang-(1-7) and PAMAMNH in complex at acidic pH.

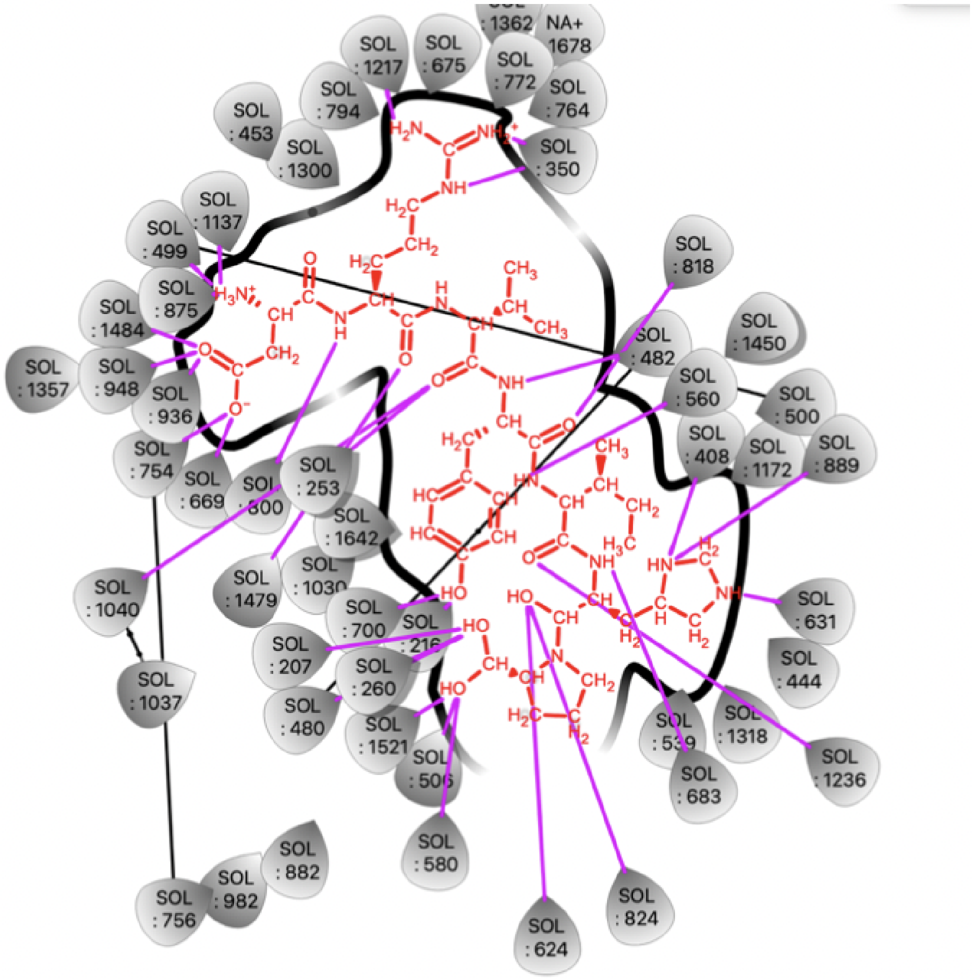


| Acceptor | Donor |
|---|---|
| WO(1137) | ASP1- $NH_3^+$ |
| WO(499) | ASP1- $NH_3^+$ |
| ASP1-(C=O) | WO(1484-948-936) |
| ASP1-($CO^-$) | WO(754-669) |
| WO (350) | ARG2-$NH_2^+$ |
| WO (1217) | ARG2-$H_2N$ |
| WO(350) | ARG2-NH |
| WO(800) | ARG2(NH) |
| ARG2-(C=O) | WO(1479) |
| VAL3(C=O) | WO(1479-1040) |
| WO(482-818) | TYR4(NH) |
| TYR4(C=O) | WO(482) |
| WO(560) | ILE5(NH) |
| ILE5(C=O) | WO(1236) |
| WO(889-408) | HIS6(HND) |
| WO(631) | HIS6(HNE) |
| WO(683) | HIS6(NH) |
| WO(624-824) | HIS6(OH) |
| WO(580-1521-506) | PRO7(HO) |

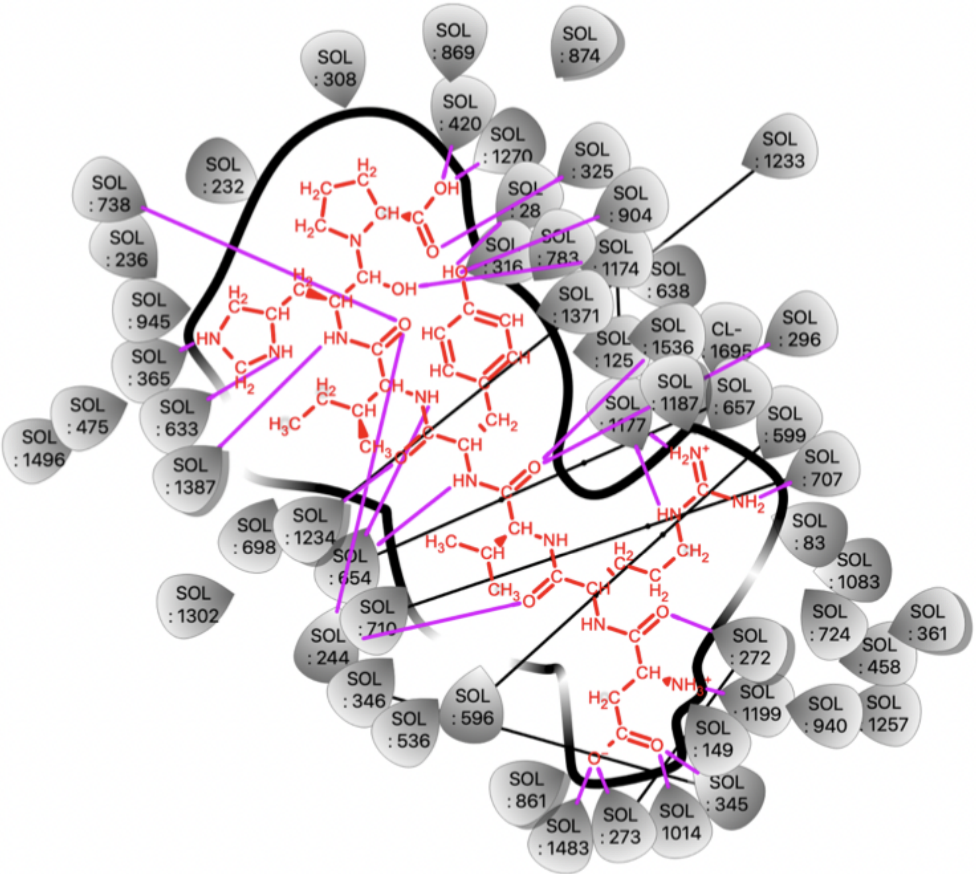


| Acceptor | Donor |
|---|---|
| ASP1(C=O) | WO(345-1014) |
| ASP1(C-O$^-$) | WO(273-1483) |
| WO(1199-272) | ASP1($NH_3^+$) |
| ASP1(C=O) | WO(272) |
| ARG2(C=O) | WO(710) |
| WO(1177) | ARG2($H_2N^+$) |
| WO(1177) | ARG2(HN) |
| VAL3(C=O) | WO(296-1536) |
| WO(904-316) | TYR4(OH) |
| ILE5(C=O) | WO(244-738) |
| HISH6(ND) | WO(633) |
| HISH6(NE) | WO(365) |
| HISH6(NH) | WO(1387) |
| WO (1270-420) | PRO7-OH |
| PRO7(C=O) | WO(325) |

Figure S16: Details on the type of hydrogen bonds that are able to be formed between Ang-(1-7) and water in complex at acidic pH.
